# Genome-wide identification and expression analysis of CDPK proteins in agarwood-producing Aquilaria agallocha trees

**DOI:** 10.1101/2025.04.27.650281

**Authors:** Khaleda Begum, Ankur Das, Raja Ahmed, Suraiya Akhtar, Sofia Banu

## Abstract

Calcium is the most common secondary messenger of the plant signal transduction mechanism. The Calcium-dependent protein kinases (*CDPK*s), a plant multi-gene family protein, act as sensors of calcium ion concentration inside plant cells. *CDPK*s convert variations in calcium concentration into a signal and phosphorylate various proteins that translate calcium signals into physiological reactions. *Aquilaria agallocha* is an aromatically important crop due to the production of valuable fragrant resinous agarwood in their heartwood. The trees produce agarwood when wounded and exposed to biotic and abiotic stresses. *CDPK*s are essential in plant development and growth, biotic and abiotic stress responses, and phytohormone-mediated signalling pathways. A comprehensive investigation of the stress-combating gene family (CDPK) in the *A*. *agallocha* genome is currently lacking. In this work, we used a bioinformatics approach to examine the entire genome of *A*. *agallocha* and identified 24 *CDPK* genes. According to the molecular phylogenetic relationships, the putative Aa*CDPK*s are grouped into four groups. Synteny analysis identified a few conserved segments (orthologous genes) between *A*. *agalloch*a and *Arabidopsis thaliana* (eight pairs), *A*. *sinensis* (twenty pairs), *Glycine max* (sixteen pairs), *Solanum tuberosum* (five pairs), and *Vitis vinifera* (ten pairs). Duplication analysis of *AaCDPK* genes indicated that dispersed duplications significantly contributed to the expansion of the *CDPK* gene family of *A*. *agallocha.* The observed AaCDPK-RBOH interaction within the protein interaction network suggests that this interaction may be crucial in integrating Ca^2+^ and ROS signaling pathways. RNA-seq data analysis shows differential expression of seven putative *AaCDPK* genes in agarwood tissue. qRT-PCR analysis revealed that six out of ten selected *AaCDPK* genes exhibited altered expression levels in response to MeJA, H_2_O_2_, and CaCl_2_ treatments. Integrated promoter analysis, PPI data, and in silico gene expression validated with qRT-PCR-analysis collectively suggest the role of CDPK-RBOH as a signalling molecule in initiating phytohormone-mediated agarwood resin formation. This study provides a strong foundation for future investigations into the study of functional roles and regulatory mechanisms of *AaCDPK* genes in sesquiterpene biosynthesis and agarwood formation.

## Introduction

Secondary messengers of plant cells provide resistance against cold, drought, heat, and high salinity by activating various physiological and metabolic responses (Valliyodan et al., 2006; Vladimir et al., 2009). Ca^2+^ is an essential secondary messenger in plants, which is critical in regulating various aspects of plant life, such as growth, development, and responses to environmental factors. (White et al., 2003; Edel et al., 2017; Wen et al., 2020. Cytosolic Ca^2+,^ acting as a secondary messenger upon modulation of its concentration by external stress, activates intracellular phosphorylation cascades. This process facilitates signal transduction, leading to the induction of physiological and biochemical alterations that enhance plant stress tolerance (Zhang et al., 2015; Ghosh et al., 2021; Li et al., 2022). Three distinct protein families primarily transduce calcium signaling in plants: CBL/CIPK complexes, Calcium-dependent protein kinases (CDPKs), and proteins belonging to the calmodulin superfamily (CaM and CML) (Chen et al., 2023; Symond et al. 2024). Although the three mentioned proteins are calcium-binding proteins critical for cellular signaling, their functional specificities arise from structural and interactional differences (Xiao et al., 2022). CaM and CMLs serve as broad-range calcium sensors. CBL proteins, however, specifically interact with CIPK kinases. CDPKs are of particular interest among these three groups of sensor proteins because they constitute a unique type of Ca^2+^ sensor and effector as they can directly convey the upstream Ca^2+^ signal to downstream signaling of protein phosphorylation (Asano et al., 2005; Hamel et al., 2014; Yip Delormel et al., 2019;). CDPK proteins are distinct due to the integration of calcium-binding and signaling properties within their single gene product (Liese and Romies, 2013; Hamel et al., 2014). CDPKs, a class of enzymes, are present in plants and some protozoa but not detected in animals and fungi (Hettenhausen et al., 2016; Sharma et al., 2021)

CDPKs possess a modular architecture comprising four distinct domains: an N-terminal variable region, a Ser/Thr kinase catalytic domain, an autoregulatory/autoinhibitory domain, and a calmodulin-like domain (Shu et al., 2020; Kumar et al., 2023). The N-terminal domains contained myristoylation and palmitoylation sites, which assisted CDPKs in anchoring or dissociating from cell membranes. The N-terminal terminal domain of CDPK proteins had the most substantial sequence divergence (Wen et al., 2020). The catalytic protein kinase domain, which has an ATP binding site, is followed by the autoinhibitory junction domain and the calmodulin-like domain, which has essential EF-hands chiral structures (1-4 in numbers) for binding to calcium (Yu et al., 2018; Mu et al., 2024). The N- and C-terminal domains of plant CDPKs exhibit distinct lengths and amino acid sequences. (Stael et al. 2011; Hettenhausen et al., 2016, Simeonovic et al., 2016). CDPK activation involves calcium-induced detachment of the junction domain from the kinase domain (Vitart et al. 2000). By phosphorylating their substrate proteins, activated CDPKs can drive Ca^2+^ signaling, which is crucial for signal transduction during abiotic stresses and microbial interactions (Schulz et al., 2013; Shu et al., 2020; Du et al., 2023).

Ca^2+^ signals orchestrate diverse plant growth and developmental processes (Kudla et al., 2010; Yuan et al., 2017). CDPKs act as Ca^2+^ sensors, regulating various aspects of plant development, including stem and petiole elongation (Matschi et al., 2013), pollen tube growth (Myers et al., 2009), and seed development (Milo Frattini, 1999). CDPKs also play crucial roles in plant stress responses (Romies et al., 2001; Asano et al., 2012). Overexpression of certain CDPKs in Arabidopsis and other plants enhances drought and salt stress tolerance (Zhu et al., 2007; Zou et al., 2010; Jiang et al., 2013; Zou et al., 2015). Beyond their established roles in stress responses, CDPKs are also critical components of plant immunity (Boudsocq and Sheen, 2013; Bredow and Monaghan, 2019. Over the past decade, advancements in plant genomic analysis have paralleled the characterization of numerous plant *CDPK*s and their functions. The *CDPK* gene was first identified as being activated by calcium ions in pea shoot membranes (Hetherington, A. M., & Trewavas, A. 1984). Members of the *CDPK* gene family have now been identified in many plant species. Exploration of *CDPK* studies showed substantial variation in the number of *CDPK* genes existing in different plant genomes. Specifically, *Arabidopsis*, *Oryza sativa*, *S*. *tuberosum*, *Zea mays*, and *Cucumis melo*, *Gossypium barbadense*, *Raphanus sativus* genome possess 34, 31, 26, 18, 40, 84 and 37 *CDPK* genes, respectively (Cheng et al., 2002; Ray et al., 2007; Fantino et al., 2017; Zhang et al., 2017, Zhao et al., 2021a; Shi., G.& Zhu., X., 2022 and Yang et al. 2023,). C*DPK* genes are emerging as pivotal players in plant defense, regulating Ca^2+-^mediated signaling pathways that are indispensable for survival. However, this gene family remains unexplored in agarwood-producing *Aquilaria* species.

The trees in the *Aquilaria* genus of the Thymelaceae plant family are unique because of the production of fragrant, resinous heartwood called agarwood (Islam and Banu, 2021. Zhang et al., 2021; Gogoi et al., 2023). Agarwood is formed in the heartwood of *Aquilaria* species in response to stresses such as physical damage (cutting), insect attacks, and infections by pathogenic microorganisms like fungi and bacteria (Chhipa and Kaushik, 2017; Gadiran et al., 2023). Agarwood production exhibits high economic viability, underpinned by a global market valued at several billion USD. This valuation is sustained by the increasing demand for a diverse product suite, encompassing wood, chips, powder, oil, dust, incense, and perfumes, all characterized by their distinctive fragrance. (Barden et al. 2000, Adhikari et al. 2021, Shivanand et al. 2022).

Intrinsically, trees in the *Aquilaria* genera produced agarwood as a self-defense mechanism (Naziz et al., 2019; Faizal et al., 2021; Gogoi et al., 2023. Stresses induce defense mechanisms in *Aquilaria* species, initiating secondary metabolite biosynthesis and subsequent accumulation of fragrant resinous substances (Kristanti et al., 2018; Tan et al., 2019; Begum et al., 2024). Chemically, agarwood resin comprises sesquiterpenes and 2-(2-phenyl ethyl) chromones (PECs), responsible for distinctive aroma (Das et al., 2023; Hou et.al., 2024). Major secondary metabolites of agarwood are synthesized via the mevalonate (MVA) and dexylulose phosphate (DPX) pathways (Xu et al., 2014; Tan et al., 2019; Lv et al., 2022; Xu et al., 2023). Despite considerable progress in elucidating the biochemical processes of agarwood formation, the precise molecular mechanisms underlying this process remain to be fully elucidated.

Wounding triggers signaling pathways, including calcium signaling, which activate *Aquilaria* species’ defense response (Kenmotsu et al., 2011; Gao et al., 2016; Tan et al., 2019; Islam et al., 2021). The Ca^2+^ signaling in the *Aquilaria* tree triggers defense responses via H_2_O_2_, ethylene (ET), jasmonic acid (JA), and salicylic acid (SA) pathways to resist stresses (Zhang et al., 2014); Lv et al., 2019; Tan et al., 2019). The phytohormones above (ET, JA, and SA) also play crucial roles in disease resistance mechanisms in model plant systems (Clarke et al., 2000). The phytohormone JA acts as a key signal molecule to initiate the biosynthesis of sesquiterpenes via H_2_O_2_ accumulation (Xu et al., 2016; Naziz et al., 2019; Faizal et al., 2021). H_2_O_2_ is a key regulator of secondary metabolite biosynthesis in plants (Ibrahim& Jaafar, 2012; Zhao et al., 2021; Fan et al., 2024), including agarwood sesquiterpene production. Under biotic/abiotic stress conditions, RBOH proteins, generating ROS (especially H_2_O_2_), are key to defense signaling in agarwood formation (Xu et al., 2013; Wang et al., 2018; Begum et al., 2024). Cytosolic Ca^2+^ levels are sensed by CDPKs, which subsequently phosphorylate RBOH proteins (Kabayshi et al., 2007; Kimura et al., 2012; Yu et al., 2024. These signaling molecules activate MYB, MYC, and WRKY transcription factors. The activation of these transcription factors, as a result of upstream signaling cascades, regulates the expression of genes of MVA and pathways, thereby influencing agarwood plant secondary metabolite biosynthesis, particularly sesquiterpene production (Tan et al., 2019). CDPKs have been demonstrated to modulate secondary metabolite biosynthesis in various plant species, including *A*. *thaliana*, *S*. *tuberosum*, *Rubia cordifolia*, and *Glycine max* (Kiselev et al., 2025). This modulation occurs through the phosphorylation of target proteins, which may include enzymes directly involved in biosynthetic pathways or transcription factors regulating gene expression. Therefore, it is reasonable to hypothesize that *CDPK*s in *Aquilaria agallocha* act as signaling molecules like other plant species. They may also regulate defense-related gene expression and subsequent sesquiterpene biosynthesis through similar phosphorylation-dependent mechanisms. Further research is required to fully elucidate the specific roles of CDPKs and their interactions within the complex regulatory network governing agarwood formation.

However, the implication of CDPK proteins in sensing Ca^2+^ ions during stress-induced agarwood formation is evident. A systematic review of peer-reviewed literature concerning CDPK genes within the *Aquilaria* species genome revealed a single publication, Xu et al. (2013), reporting on the role of CDPK genes in agarwood formation. Transcriptome analysis of MeJA-treated *A*. *sinensis* calli (Xu et al., 2013) revealed 25 CDPK unigenes linked to Ca^2+^ signaling.

Subsequent expression profiling of five CDPK genes selected from this dataset indicated their probable involvement in agarwood deposition. previous research identified CDPK genes within the MeJA-treated transcriptome of *A. sinensis*, a comprehensive, genome-wide characterization of this gene family within the genus *Aquilaria* remains lacking. Therefore, the present study aims to identify and characterize the CDPK gene family in the *A. agallocha* genome."

## 2. Materials and Methods

### 2.1. Establishment of tissue culture for stress treatment

For tissue culture, leaves of *A. agallocha* were cleaned with sterile water and sterilized with 0.1% mercuric chloride solution. After that, the leaves were divided into small pieces and aseptically inoculated in Murashige and Skoog (MS) medium (pH 5.8) with 3% (w/v) sucrose enriched with 2, 4-dichlorophenoxyacetic acid (2, 4-D) at 6 mg/L and kinetin at 2 mg/L, then it was incubated at 25 °C in the dark for an extended period (Okudera et al. 2009). Suspension cultures were established for stress treatments. Healthy, friable calli were treated with 0.1 mM MeJA, 0.1 mM H₂O₂, and 10 mM CaCl₂ in suspension culture at 130 rpm for 5 days (Kumeta and Ito 2010). The calli without any stress treatment is considered as a control samples. The samples were harvested at zero h, one h, three h, six h, 24 h, and 48 h after treatment.

### 2.2. Retrieval and identification of *CDPK* genes in the genome of *Aquilaria agallocha*

To systematically identify the *CDPK* gene family in the genome of *Aquilaria agallocha,* we performed the following steps: Step 1. We collected data from genomic sequences from an earlier annotation project (Das et al., 2021). Step 2, The *CDPK* structural-specific protein kinase domain (PF00069) and EF-hand domain (PF13499) sequences alignment were retrieved from the Pfam database (http://pfam.xfam.org/browse); step 3, The obtained alignments were used to build hidden Markov model (HMM), and then hmmsearch program was utilized to identify the putative CDPK proteins with the HMMER 3.3.2 software (Potter et al., 2018); step 4. The resultant short and duplicate sequences were removed after screening, step 5. The remaining deduced sequences were then analyzed to confirm the presence of those mentioned two conserved domains in the SMART software tool (Simple Modular Architecture Research Tool; (http://smart.embl-heidelberg.de/) and NCBI conserved domain database (https://www.ncbi.nlm.nih.gov/Structure/cdd/cdd.shtml) and step 6. The obtained protein sequences of A. *agallocha* putative considered CDPK and were named based on homology with the Arabidopsis CDPK proteins (Li et al., 2022). The identification of CDPK proteins in the Aquilaria genome was also performed using Blast2-Go software. Using Blast2GO, a BLASTP search of 25,234 A. agallocha proteins was performed against the SwissProt database.

### 2.3. Analysis of Physiological Properties and prediction of sub-cellular localization

The physiological properties of the identified 24 putative *AaCDPK* were analyzed with the ExPASy-ProtParam tool (available online: http://web.expasy.org/protparam/) (Wen et al., 2020). The N-terminal myristoylation sites of putative *AaCDPKs* were predicted using the ExpasyMyristoylator program online tool (https://web.expasy.org/myristoylator/), and palmitoylation site was predicted by CSS-plam4.0 (http://csspalm.biocuckoo.org/). The subcellular location of the putative AaCDPKs was expected with the online protein subcellular location predictor WoLF PSORT (https://wolfpsort.hgc.jp/).

### 2.4. Phylogenetic relationship of *AaCDPK*

To analyze the evolutionary relationship AaCDPK with the AtCPK, the CPK protein sequences of *A*. *thaliana* were downloaded from The Arabidopsis Information Resource (TAIR, https://www.arabidopsis.org/) and NCBI. Multiple sequence alignment of AaCDPKs and AtCPKs was performed with the ClustalW program. The Phylogenetic tree was prepared using the obtained MSA, following the neighbor-joining method with 1000 bootstrap values utilizing the p-distance method of MEGA-X. The displayed phylogenetic tree was constructed by uploading the Newick format of MEGA-X to iTOL online website (https://itol.embl.de/).

### 2.5. Analysis of gene and protein motif structure, Conserved residue of AaCDPK

The split gene structure of *AaCDPKs* was revealed with the Gene Display Server (http://gsds.cbi.pku.edu.cn)) by aligning *AaCDPK* CDS sequences with genomic sequences of *AaCDPK*s. The motif structure of *AaCDPK* proteins was analyzed using MEME suits software. The default parameters of the MEME suite were followed except for the studied number of motifs set as ten to illustrate the motif pattern of *AaCDPK* proteins (https://meme-suite.org/meme/).

### 2.6. Cis-Regulatory elements and analysis of in-silico gene Expression

The Cis-regulatory elements in the promoter region of *AaCDPK*s were analyzed using the PlantCARE database (http://bioinformatics.psb.ugent.be/webtools/plantcare/html/). The 2000 bp upstream of the positive (+) strand located *AaCDPK* genes and the downstream 2000 bp of negative (–) strand located *AaCDPK* genes were considered to identify the Cis Regulatory elements.

To investigate the expression of *AaCDPK* transcript in infected and healthy trees of *A. agallocha*, raw RNA-seq data in FASTQ format (Accession number PRJNA449813 ID: 449813, SRX4184708-SRX4184710, and SRX4149019-SRX4149021) were downloaded from the NCBI SRA database (https://www.ncbi.nlm.nih.gov/sra). The differentially expressed transcript was calculated using the HISAT2 StringTie-DESeq2 workflow (Kim et al., 2015; Love et al., 2014; Kovaka et al., 2019).

### 2.7. Gene Ontology (GO) annotation

Functional annotation of potential AaCDPK proteins was performed using OmicsBox software (https://www.blast2go.com/). The amino acid sequences of putative AaCDPK were imported into the Blast2Go software. We performed the mentioned three steps to annotate the putative AaCDPK proteins: 1) BlastP against the Swissport protein database, 2) mapping and retrieval of GO-term linked with BLASp findings, and 3) annotation of GO terms connected with each query sequence to know the protein function (Lopez-Ortis et al., 2019, Vishwakarma et al., 2024).

### 2.8. Protein-protein interaction

STRING v 12 software (https://string-db.org/) was used to determine protein-protein interactions. All AaCDPK protein sequences were uploaded, and "*Arabidopsis thaliana*" was selected from the list of organisms. The interaction network was determined based on text mining, databases, coexpression, neighborhood, gene fusion, co-occurrence, and experimental evidence. The resultant PPI images were saved as.jpeg picture files.

### 2.8 Synteny and duplication events

The Multiple Collinearity Scan toolkit (MCScanX) was employed to evaluate the syntenic relationships between *A*. *agallocha*- *A*. *thaliana*, *A. agallocha*- *Solanum tuberosum*, *A*.*agallocha*- *Glycine max* and *A. agallocha*-*Vitis venifera*. The Dual Synteny Plott function of TBtools-II (Chen et al., 2023) was used to construct the graphs for the syntenic relationship analysis (Bettaieb and Bouktila 2020). The Advanced Circos view program of TBtools II was utilized to create a map illustrating the syntenic relationship between the orthologous *AaCDPK* genes of the Agarwood plant. (Sami et al., 2024). The pattern of AaCDPK gene duplication was found with the help of Dupgen finder (https://github.com/qiao-xin/DupGen_finder). Subsequently, the synonymous substitution rate (Ks), non-synonymous substitution rate (Ka), and the Ka/Ks ratio between the duplicated *AaCDPK* gene pairs were calculated using the KaKs calculator of TBtools II (Chen et al., 2023). Evolutionary divergence time (Millions of Years, MLA) was measured using a well-known formula, T = Ks/2λ x 10^−6^, where λ = 6.5 × 10^−9^ (Sun et al., 2022).

### 2.9. Homology modelling

Homology modeling was employed to determine the 3D structures of the AaCDPK proteins using the Swiss Model web server (https://swissmodel.expasy.org/). Automated protein structure homology-modeling server, swiss-model used for generating 3D models of AaCDPKs (https://swissmodel.expasy.org). The structural assessment of (validation and correctness) of homology-modelled protein was assessed based on MolProbity Score, Clash Score, and Ramachandran favored % (https://swissmodel.expasy.org/assess). The quality of the predicted 3D protein structures was evaluated using the ERRAT server within the Structural Analysis and Verification Server (SAVES) v6.0. (https://saves.mbi.ucla.edu/). Projected models of AaCDPK proteins were extracted in various 3D locations using UCSF CHIMERA 1.10. (Mu et al., 2024)

### 2.10. RNA extraction, cDNA conversion, and RT-qPCR

Total RNA was extracted from *A*. *agallocha* calli tissue using the RNeasy Plant Mini Kit (Qiagen) according to the manufacturer’s instructions. The quality of extracted RNA was determined on 1% agarose gel electrophoresis by visualization and quantified with the Multiskan Sky Microplate Spectrophotometer of Thermo Fisher Scientific, USA. One microgram of RNA was used to synthesize the first strand of cDNA with SuperScript III Reverse Transcriptase (Thermofisher). The qRT-PCR was conducted using a QuantStudio™ 3 real-time PCR system (Applied Biosystems, USA) and the PowerUp SYBR Green Master Mix (AppliedBiosystems).

Ten *AaCDPK*s gene-specific primer pairs were designed with IDT PrimerQuest software (https://sg.idtdna.com/pages/tools/primerquest/) and are enlisted in **Supplementary Table 1**. The standardized GAPDH primer was utilized as the internal control (Islam et al., 2020). For each biological replicate, the analyses were performed with three technical replicates, each containing 20 μl of reaction volume in optical stripes, the temperature pattern of 95°C for 1 min, followed by 40 cycles at 95°C for 10 s and 60°C for 30 s, was followed as thermal cycler profile. Fold change in the gene expression was measured by the delta-delta CT method (Ding et al., 2021)

### 2.11. Statistical analysis

The library of RNA-seq data was normalized using the median ratios (Andres and Huber 2010) with p < 0.05. Log fold change [1, and\- 1 were considered differentially upregulated and down-regulated, respectively. Data of ct value are presented as the mean ± standard error of the mean from three biological replicates. Asterisks indicate statistically significant differences compared to the control group (1) as determined by Student’s t-test analysis (*P ≤ 0.05; P ≤ 0.01) (Yan et al., 2018).

## 3. Results

### 3.1. Identification of *CDPK* genes from *A. agallocha* plants

A total of 24 *CDPK* family genes were identified in the *A. agallocha* genome by local BLASTp and HMM search. Using Blast2GO, a BLASTP search identified homologs of the query sequences based on sequence similarity. Selecting sequences containing the protein kinase domain (IPR000719) resulted in 649 sequences. An additional filtering step of targeting the EF-hand domain (IPR002048) resulted in 38 sequences. After SMART search and Interpro scan search, A final set of 24 protein sequences was identified as *A. agallocha* CDPK protein, based on the presence of two characteristic domains (the protein kinase domain and the EF-hand domains) of plant CDPK protein. The HMMER analysis, targeting the PF00069 and PF13499 domains, yielded similar results with the Blast2Go.

### 3.2. Physiological properties of *AaCDPK*s

The physiological properties of AaCDPK, including CDS length, peptide length, number of conserved EF-hands, pI, molecular weight, myristoylation sites, and palmitoylation sites, were analyzed using Expasy-ProtParam and are presented in **Table 1**. The *AaCDPK*s identified in our investigation varied significantly, ranging from 321 amino acids (AaCDPK24.2) to 606 amino acids (*AaCDPK10*), the molecular weights (MW) ranged from 36.676998 KDa (*AaCDPK24.2*) to 68.12337 KDa (AaCDPK*10*) and their predicted isoelectric (PI) points ranged from 5.13 (*AaCDPK20*, *AaCDPK24.2*) to 9.23 (*AaCDPK16*). Subcellular localization analysis predicted that most proteins were localized to the cytoplasm, while a smaller proportion was predicted to reside in chloroplasts, mitochondria, the ER membrane, peroxisomes, and the cytoskeleton. Most AaCDPK genes have three or four numbers of EF-hand for calcium sensing, except AaCDPK13.2, which has two EF-hand domains. Among 24 *AaCDPK*, myristoylation sites were present in nine putative *AaCDPK*s, and palmitoylation sites were present in all the putative *AaCDPK*s. An equal distribution of AaCDPKs was observed between the plus and minus strands (50% each) (**Supplementary Table 2**).

**Table 1:**
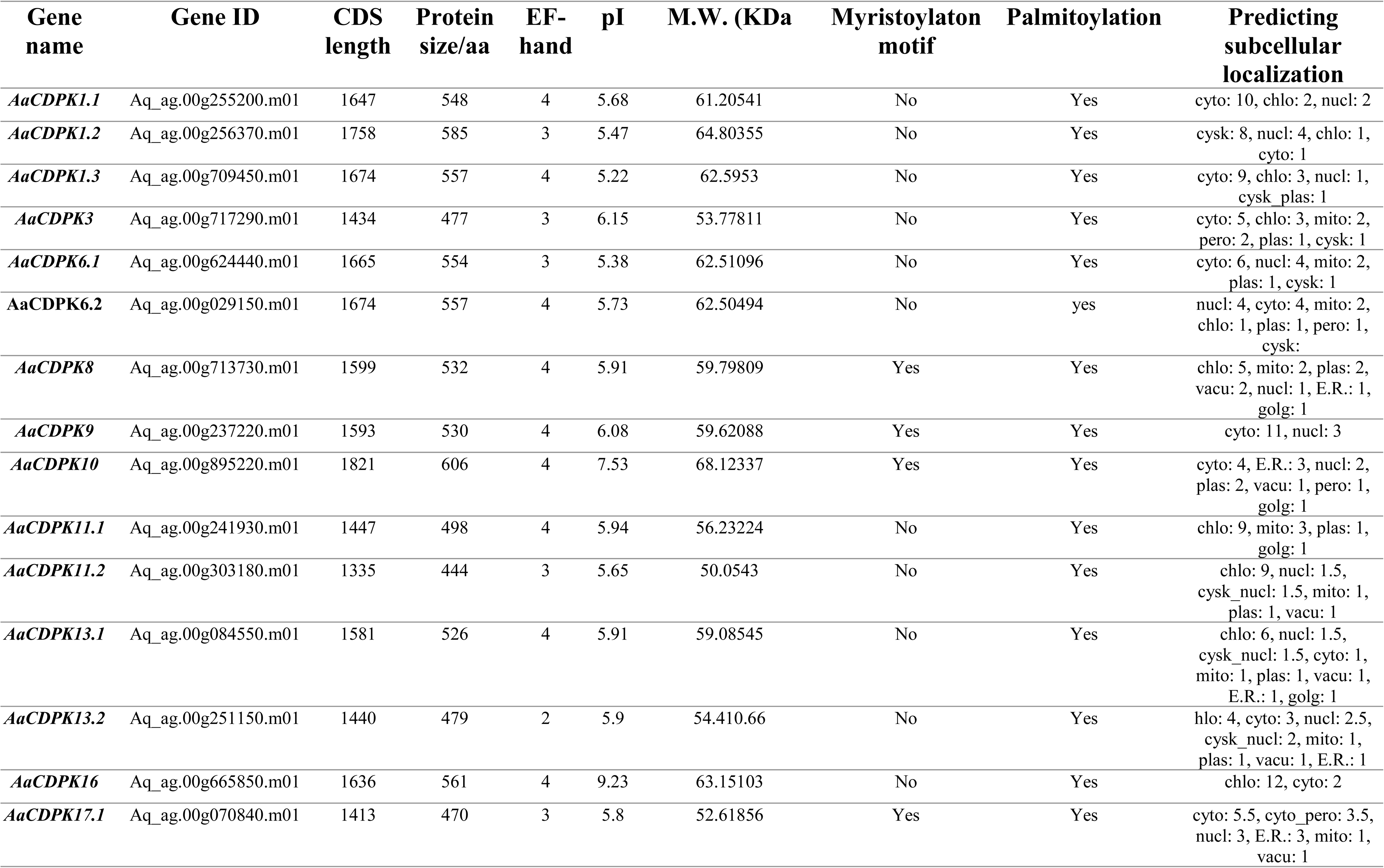
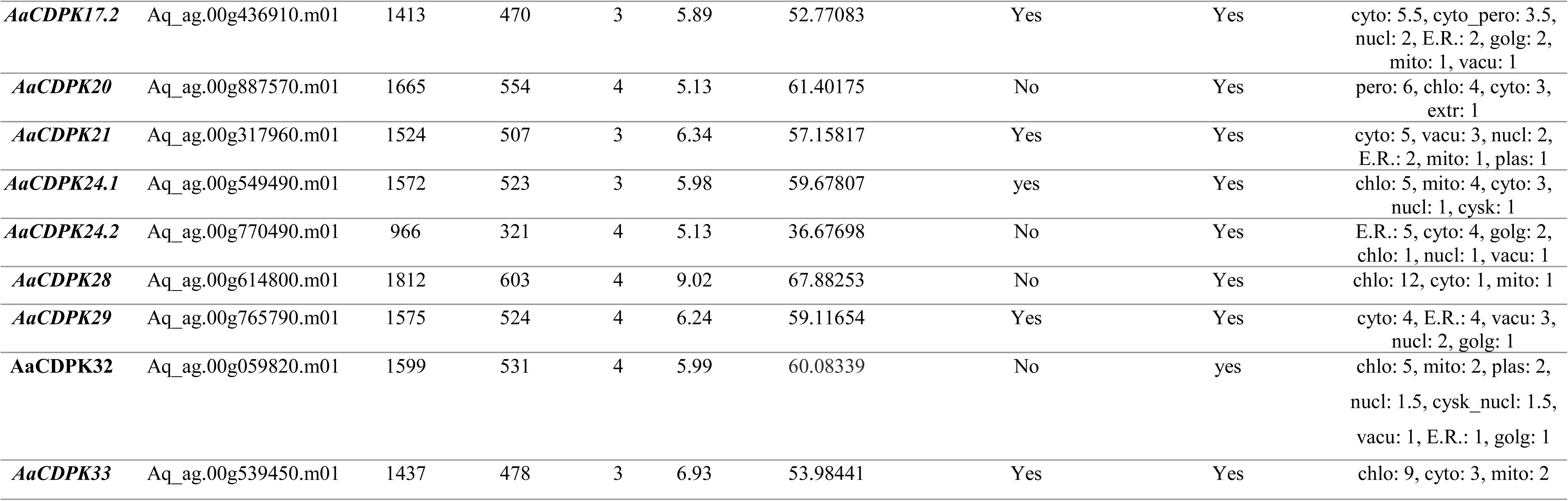
Details of the 24 putative AaCDPK genes identified in this study.

| Gene name | Gene ID | CDS length | Protein size/aa | EF-hand | pI | M.W. (KDa | Myristoylaton motif | Palmitoylation | Predicting subcellular localization |
| --- | --- | --- | --- | --- | --- | --- | --- | --- | --- |
| *AaCDPK1.1* | Aq_ag.00g255200.m01 | 1647 | 548 | 4 | 5.68 | 61.20541 | No | Yes | cyto: 10, chlo: 2, nucl: 2 |
| *AaCDPK1.2* | Aq_ag.00g256370.m01 | 1758 | 585 | 3 | 5.47 | 64.80355 | No | Yes | cysk: 8, nucl: 4, chlo: 1, cyto: 1 |
| *AaCDPK1.3* | Aq_ag.00g709450.m01 | 1674 | 557 | 4 | 5.22 | 62.5953 | No | Yes | cyto: 9, chlo: 3, nucl: 1, cysk_plas: 1 |
| *AaCDPK3* | Aq_ag.00g717290.m01 | 1434 | 477 | 3 | 6.15 | 53.77811 | No | Yes | cyto: 5, chlo: 3, mito: 2, pero: 2, plas: 1, cysk: 1 |
| *AaCDPK6.1* | Aq_ag.00g624440.m01 | 1665 | 554 | 3 | 5.38 | 62.51096 | No | Yes | cyto: 6, nucl: 4, mito: 2, plas: 1, cysk: 1 |
| *AaCDPK6.2* | Aq_ag.00g029150.m01 | 1674 | 557 | 4 | 5.73 | 62.50494 | No | yes | nucl: 4, cyto: 4, mito: 2, chlo: 1, plas: 1, pero: 1, cysk: |
| *AaCDPK8* | Aq_ag.00g713730.m01 | 1599 | 532 | 4 | 5.91 | 59.79809 | Yes | Yes | chlo: 5, mito: 2, plas: 2, vacu: 2, nucl: 1, E.R.: 1, golg: 1 |
| *AaCDPK9* | Aq_ag.00g237220.m01 | 1593 | 530 | 4 | 6.08 | 59.62088 | Yes | Yes | cyto: 11, nucl: 3 |
| *AaCDPK10* | Aq_ag.00g895220.m01 | 1821 | 606 | 4 | 7.53 | 68.12337 | Yes | Yes | cyto: 4, E.R.: 3, nucl: 2, plas: 2, vacu: 1, pero: 1, golg: 1 |
| *AaCDPK11.1* | Aq_ag.00g241930.m01 | 1447 | 498 | 4 | 5.94 | 56.23224 | No | Yes | chlo: 9, mito: 3, plas: 1, golg: 1 |
| *AaCDPK11.2* | Aq_ag.00g303180.m01 | 1335 | 444 | 3 | 5.65 | 50.0543 | No | Yes | chlo: 9, nucl: 1.5, cysk_nucl: 1.5, mito: 1, plas: 1, vacu: 1 |
| *AaCDPK13.1* | Aq_ag.00g084550.m01 | 1581 | 526 | 4 | 5.91 | 59.08545 | No | Yes | chlo: 6, nucl: 1.5, cysk_nucl: 1.5, cyto: 1, mito: 1, plas: 1, vacu: 1, E.R.: 1, golg: 1 |
| *AaCDPK13.2* | Aq_ag.00g251150.m01 | 1440 | 479 | 2 | 5.9 | 54.410.66 | No | Yes | hlo: 4, cyto: 3, nucl: 2.5, cysk_nucl: 2, mito: 1, plas: 1, vacu: 1, E.R.: 1 |
| *AaCDPK16* | Aq_ag.00g665850.m01 | 1636 | 561 | 4 | 9.23 | 63.15103 | No | Yes | chlo: 12, cyto: 2 |
| *AaCDPK17.1* | Aq_ag.00g070840.m01 | 1413 | 470 | 3 | 5.8 | 52.61856 | Yes | Yes | cyto: 5.5, cyto_pero: 3.5, nucl: 3, E.R.: 3, mito: 1, vacu: 1 |
| *AaCDPK17.2* | Aq_ag.00g436910.m01 | 1413 | 470 | 3 | 5.89 | 52.77083 | Yes | Yes | cyto: 5.5, cyto_pero: 3.5, nucl: 2, E.R.: 2, golg: 2, mito: 1, vacu: 1 |
| *AaCDPK20* | Aq_ag.00g887570.m01 | 1665 | 554 | 4 | 5.13 | 61.40175 | No | Yes | pero: 6, chlo: 4, cyto: 3, extr: 1 |
| *AaCDPK21* | Aq_ag.00g317960.m01 | 1524 | 507 | 3 | 6.34 | 57.15817 | Yes | Yes | cyto: 5, vacu: 3, nucl: 2, E.R.: 2, mito: 1, plas: 1 |
| *AaCDPK24.1* | Aq_ag.00g549490.m01 | 1572 | 523 | 3 | 5.98 | 59.67807 | yes | Yes | chlo: 5, mito: 4, cyto: 3, nucl: 1, cysk: 1 |
| *AaCDPK24.2* | Aq_ag.00g770490.m01 | 966 | 321 | 4 | 5.13 | 36.67698 | No | Yes | E.R.: 5, cyto: 4, golg: 2, chlo: 1, nucl: 1, vacu: 1 |
| *AaCDPK28* | Aq_ag.00g614800.m01 | 1812 | 603 | 4 | 9.02 | 67.88253 | No | Yes | chlo: 12, cyto: 1, mito: 1 |
| *AaCDPK29* | Aq_ag.00g765790.m01 | 1575 | 524 | 4 | 6.24 | 59.11654 | Yes | Yes | cyto: 4, E.R.: 4, vacu: 3, nucl: 2, golg: 1 |
| *AaCDPK32* | Aq_ag.00g059820.m01 | 1599 | 531 | 4 | 5.99 | 60.08339 | No | yes | chlo: 5, mito: 2, plas: 2, nucl: 1.5, cysk_nucl: 1.5, vacu: 1, E.R.: 1, golg: 1 |
| *AaCDPK33* | Aq_ag.00g539450.m01 | 1437 | 478 | 3 | 6.93 | 53.98441 | Yes | Yes | chlo: 9, cyto: 3, mito: 2 |

### 3.3. Phylogenetic Relationship of CDPKs of *A*.*agallocha* with *A*. *thaliana CDPK* genes

To study the evolutionary relationship of CDPKs in *A. agallocha* and *Arabidopsis*, we constructed a phylogenetic tree for 58 CDPKs (i.e.,24 in *A. agallocha* and 34 in *Arabidopsis*) (**Figure 1**). All CDPKs were unevenly distributed and divided into four subgroups (groups I to IV) according to the evolutionary distance. The phylogenetic tree grouped the protein sequences of all CDPK into four clusters. The phylogenetic grouping of proteins facilitated the identification of *A*. *agallocha* orthologs corresponding to those in Arabidopsis. The specific distribution of CDPKs was as follows (total: *A. agallocha* Arabidopsis): Group I (17: 7, 10); Group II (20: 7, 13); Group III (14: 6, 8); Group IV (5:2, 3) (**Figure 1**).

**Fig. 1:**
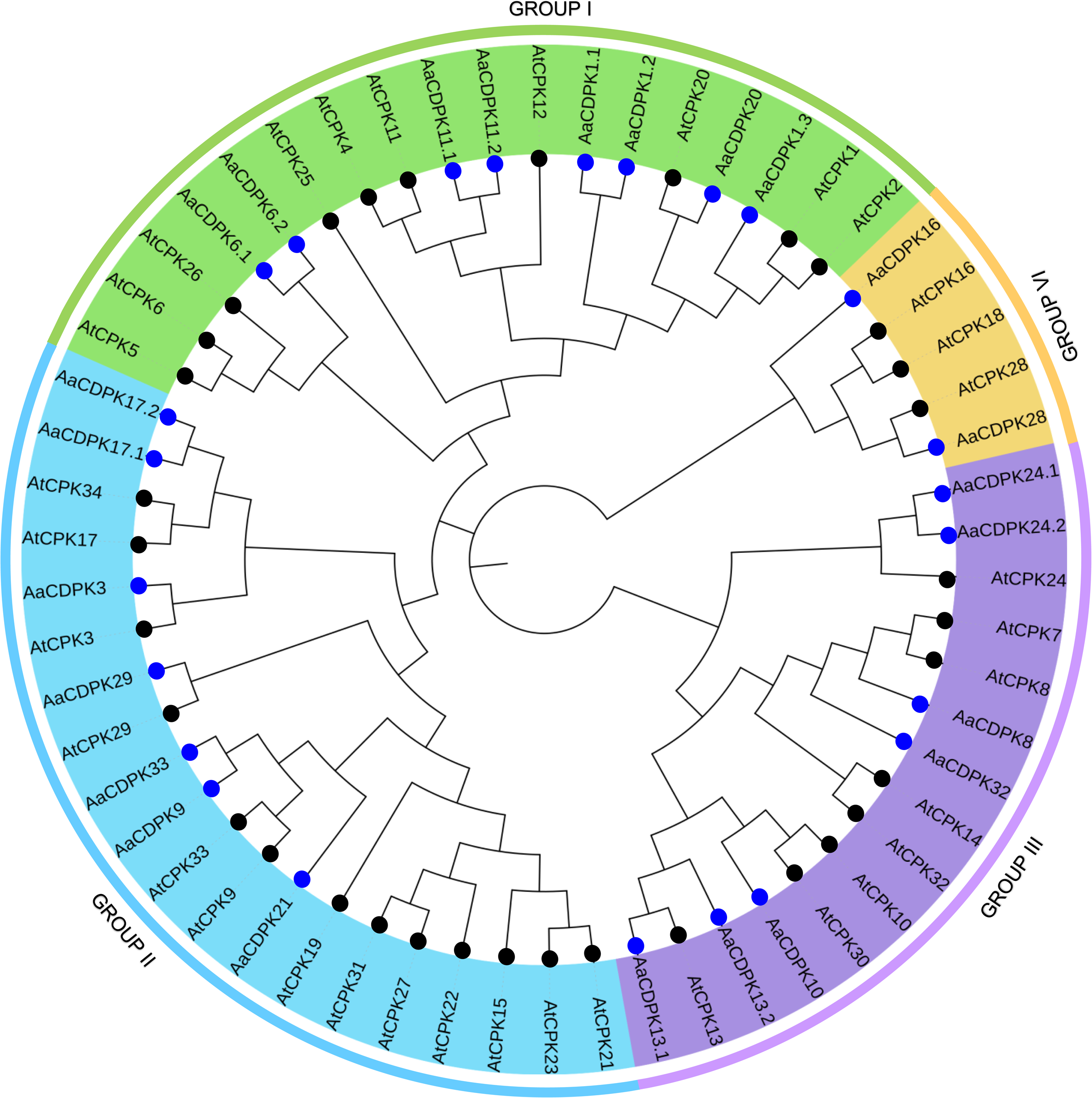
A phylogenetic tree of CDPK proteins from A. agallocha and A. thaliana. A phylogenetic tree was created using MEGA-X software employing the neighbor-joining method with 1000 bootstrap replicates. Four group (Groups 1-4) are represented by distinct color. Species abbreviations: Aa (A. agallocha) and At (Arabidopsis). The blue and black circle represent A. agallocha and Arabidopsis CDPK genes, respectively

AtCDPK2/5/6/7/12/14/15/18/19/22/23/25/26/27/30/31/32/34) from Arabidopsis have no homologous protein in *A*. *agallocha*. We speculated that significant differences in the number and type of CDPKs divided into four subgroups between *Arabidopsis* and *A. agallocha* were due to species specificity

### 3.4 Gene structures, motif composition, conserved residue

Exon-intron organization patterns were illustrated based on their evolutionary relationships to disclose the structural and functional relationships between *CDPK* genes in the genome of *A*. *agallocha*. As shown in (**Figure 2**), each Aa*CDPK* had six to thirteen number of exons. Analysis of exon counts within the CDPK protein sequences revealed a distinct pattern of variation across the four identified groups. Group IV CDPKs demonstrated the highest exon number, with AaCDPK28 and AaCDPK16 exhibiting 13 and 12 exons, respectively. A comparative analysis of the other groups indicated that two members of Group I (AaCDPK6.1 and AaCDPK6.2) and two members of Group III (AaCDPK10 and AaCDPK13.2) possessed the following highest exon counts. Subsequently, an exon count of eight was observed in two members of Group I (AaCDPK1.2 and AaCDPK11.2), two members of Group III (AaCDPK8 and AaCDPK32), and three members of Group II (AaCDPK3, AaCDPK21, and AaCDPK29). Finally, the lowest exon number, six, was identified in two members of Group III (AaCDPK24.1 and AaCDPK24.2). The findings showed that *AaCDPK* genes with more homogeneous sequences typically contain equal exons. Each CDPK differed in terms of the number and types of introns. There were 77 phase 1 introns, the most prevalent, followed by 73 phase 0 and 17 phase 2 introns. Phase 0 introns ranged from two to seven in each member (except for AaCDPK11.1, which lacks phase 0 introns), while phase 1 introns varied from two to seven in number, and phase 2 introns were present on 50% of the putative *AaCDPKs*.

**Fig. 2:**
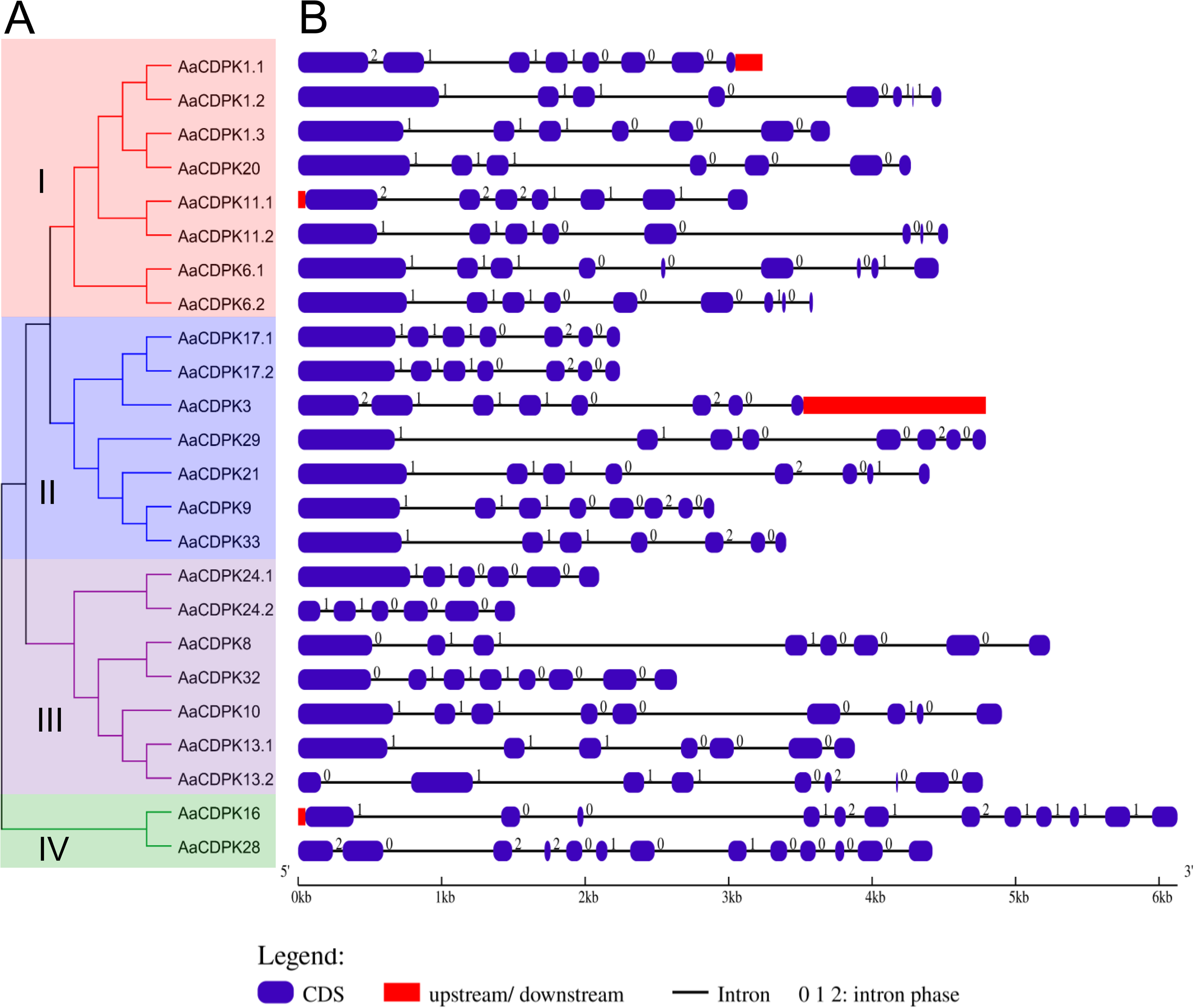
The gene structure of AaCDPKs was analyzed. (A) Phylogenetic relationships among the AaCDPK proteins were visualized using an unrooted tree, four group coded with different color. (A) The exon-intron organization of each AaCDPK gene was illustrated, where boxes represent exons and lines represent introns.

The results of MEME suites demonstrated that the motif composition of AaCDPK is very similar; most AaCDPK have five protein kinase domains and two EF-Hand domains (**Figure 4**). A member of Group III (AaCDPK24.2) contains three protein kinase domains. Motif 9 (the EF-hand domain) was not detected in three Group I members, five Group II, and one member of Group III. Furthermore, Motif 5 (also an EF-hand domain) was absent in two members of Group IV. As shown by the motif analysis in (**Figure 3**), half of the identified AaCDPKs had all the examined conserved motifs. Eleven AaCDPKs had nine motifs, while only one (AaCDPK24.2) had seven (**Supplementary Table 3).**

**Fig. 3:**
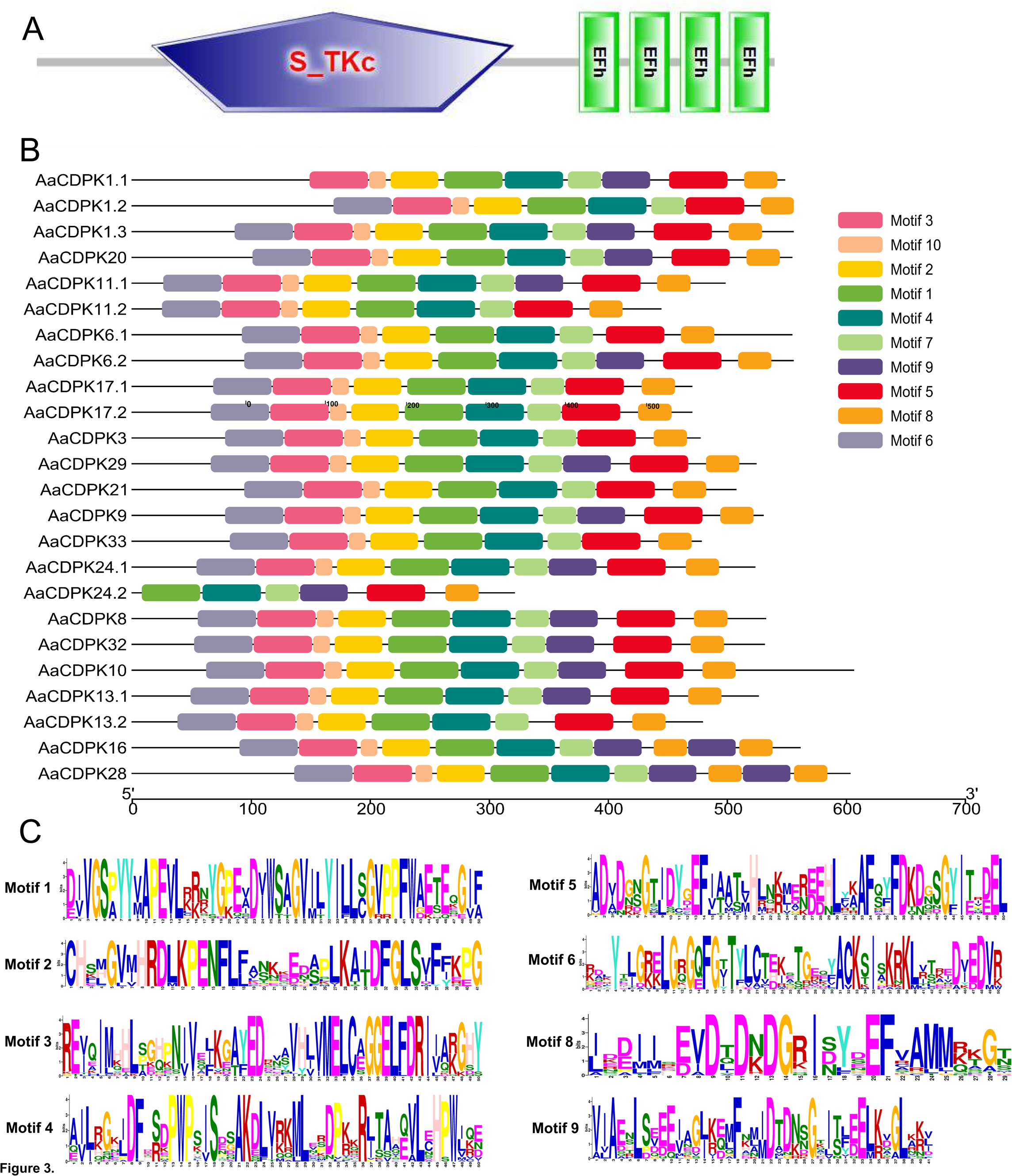
Protein structure of AaCDPKs. (A) Structure of a typical according to SMART software tool (http://smart.embl-heidelberg.de/). (B) Ten different motifs of AaCDPKs represented by different colour. (C)Sequences of ten motif. The quantity of information included at each place in the sequence is indicated by the bit score. A correlation exists between the height of each character and the degree of amino acid conservation across all proteins studied.

**Fig. 4:**
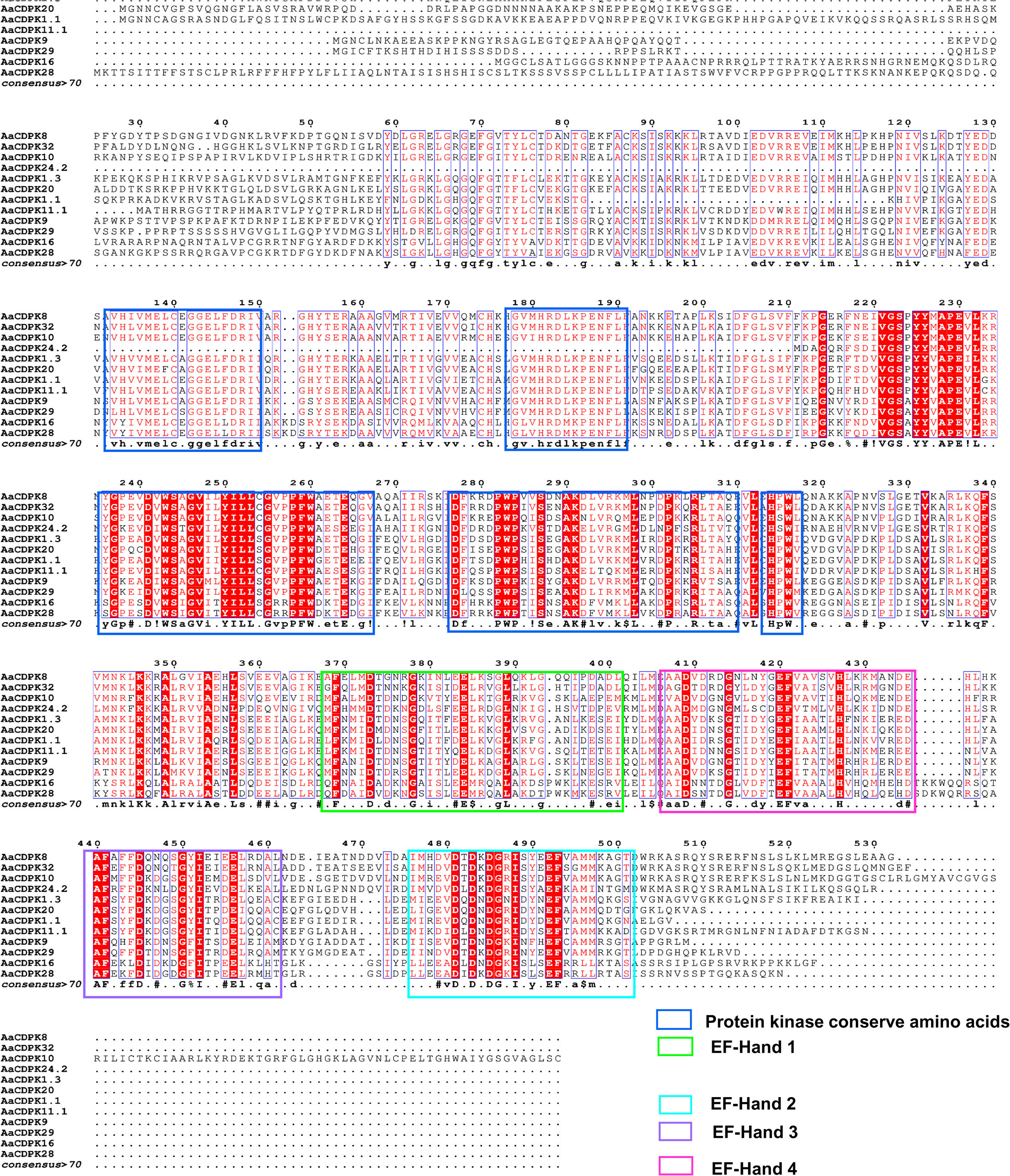
The AaCDPK protein conserved domain prepared with multiple protein sequence alignment of A. agallocha. Low levels of amino acids are represented by lighter shading, while highly conserved amino acids are indicated by red shading. Protein kinase ((PF00069) domain and EF-hand domain display with blue and four EF-hand domain (PF13499) display with four different colored boxes.

### 3.5. Promoter cis-regulatory elements and in-silico expression of AaCDPKs in infected agarwood tissue

A comprehensive analysis of AaCDPK promoter sequences demonstrated the presence of a diverse repertoire of cis-regulatory elements **(Figure 5).** These elements were subsequently classified into four distinct groups, reflecting their functional characteristics. Stress-responsive cis-regulatory elements are (ARE, DRE, DRE_Core, MBS, LTR, STRE GT1-motif, WUNmotif, ARE, WRE3, and TC-rich repeat); hormone-responsive (ABRE, ERE, AUXRR_Core, AuxRE, ERE, Pbox, GARE -motif, TCA element, TGACG-motif, CGTCA motif, and TATC-box), and plant development-related (CCAAAT-box, CAT-box, GCN4_motif, HD-Zip 1, O2-site, MBSi and A box) and core elements (AT-rich-elements, AT-rich sequences, AT-TATA-Box, CAAT-box, HD-Zip 3, TATA, TATA-Box, W box and MRE) (**Supplementary Table 4**).

**Fig. 5:**
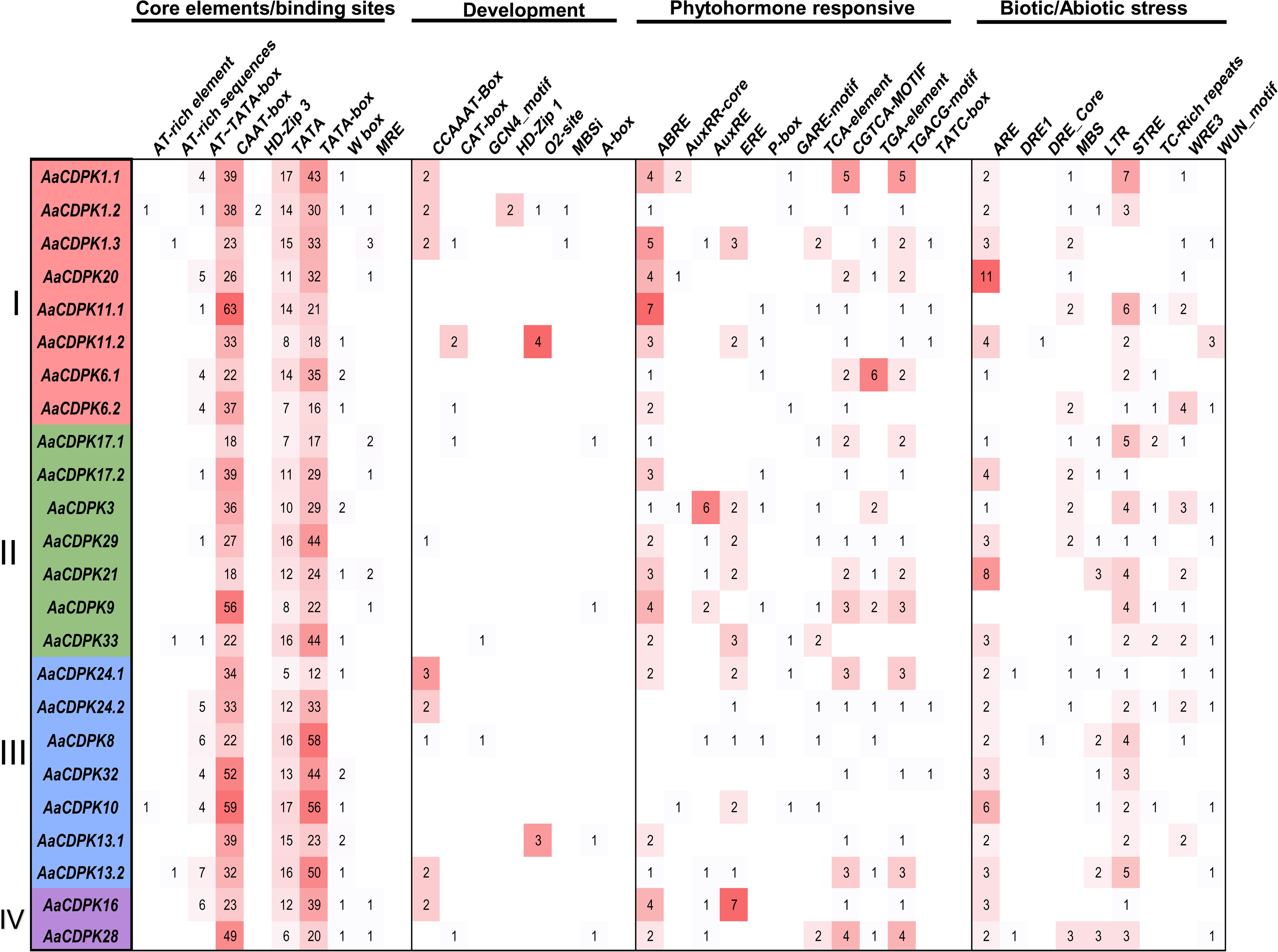
Analysis of Cis-Regulatory Elements in AaCDPK promoter. Cis-regulatory elements, of AaCDPKs categorized into four distinct groups. The intensity of red color in a visualization corresponds to the number of these cis-elements located 2000bp of the upstream or down stream of AaCDPKs

Analysis of agarwood tissue’s RNA-Seq data revealed that seven putative AaCDPKs were differentially expressed. Total six putative AaCDPKs, i.e., two members of each of the two groups—Group I (*AaCDPK1.1*, *AaCDPK20*); Group III (*AaCDPK10*, *AaCDPK32*) one member of Group II (*AaCDPK21*); —showed enhanced expression up to 1-2-fold. However, one member of two Groups, i.e., Group I (*AaCDPK1.2*) and Group II (*AaCDPK17.1*), showed up to a six-fold downregulation (**Figure 6, Supplementary Table 5**).

**Fig. 6:**
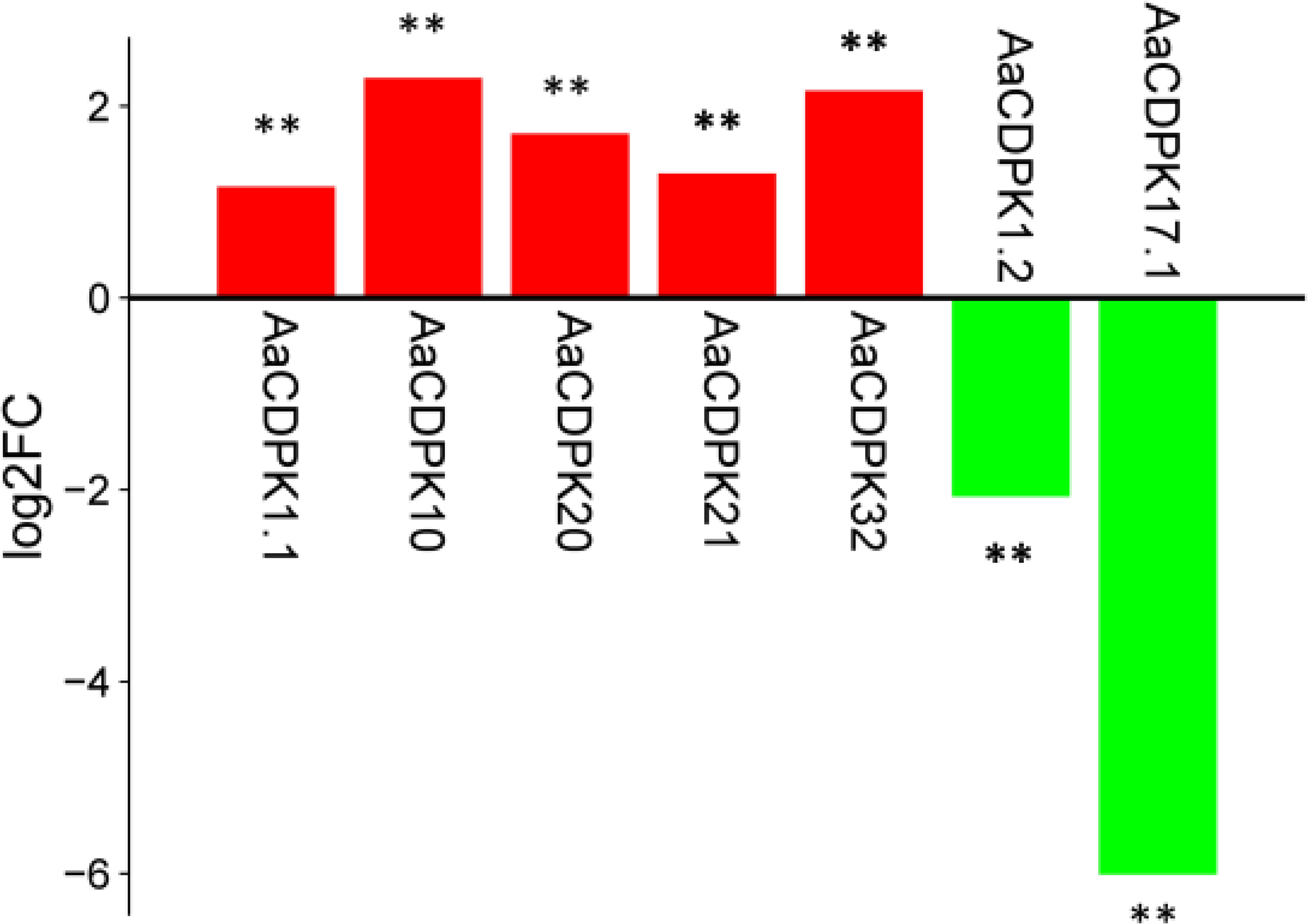
Differentially expressed AaCDPK genes in A. agallocha under stress conditions, relative to healthy controls, are illustrated in a bar diagram. The x-axis represents gene names, and the y-axis denotes the log2 fold change in expression. Data were obtained from publicly available RNA-seq data (accession number PRJNA449813)

### 3.6. Functional categorization

A Gene Ontology (GO) analysis was performed to predict the conserved functional attributes of CDPK genes across the investigated species. The results conclusively demonstrated that the putative CDPK proteins under investigation were associated with biological, cellular, and molecular functions. All putative AaCDPKs exhibited functions related to intracellular signal transduction. Other predicted biological functions include regulation of pollen tube growth, regulation of inward rectifier potassium channel activity, positive regulation of salt stress response, protein autophosphorylation, response to water deprivation and cold, regulation of anion channel activity, abscisic acid-activated signaling pathway activity, pollen maturation, negative regulation of salt stress response, positive regulation of water deprivation response, defense response to fungus, positive regulation of the abscisic acid-activated signaling pathway, response to salt stress, heat acclimation, positive regulation of the gibberellic acid-mediated signaling pathway, response to calcium ions, negative regulation of defense response to bacterium, and regulation of secondary shoot formation. The cellular functions or their cellular position inside the cell are plasma membrane, nucleus, cytoplasm, endoplasmic reticulum membrane, cytosol, etc. The predicted molecular functions of the AaCDPKs encompass calmodulin-dependent protein kinase activity, calmodulin binding, protein serine kinase activity, ATP binding, calcium-dependent protein serine/threonine kinase activity, calcium diacylglycerol-dependent serine/threonine kinase activity, protein phosphatase binding, protein serine/threonine phosphatase activity, protein tyrosine kinase activity, and metal ion binding. (**Figure 7, Supplementary Table 6**).

**Fig. 7:**
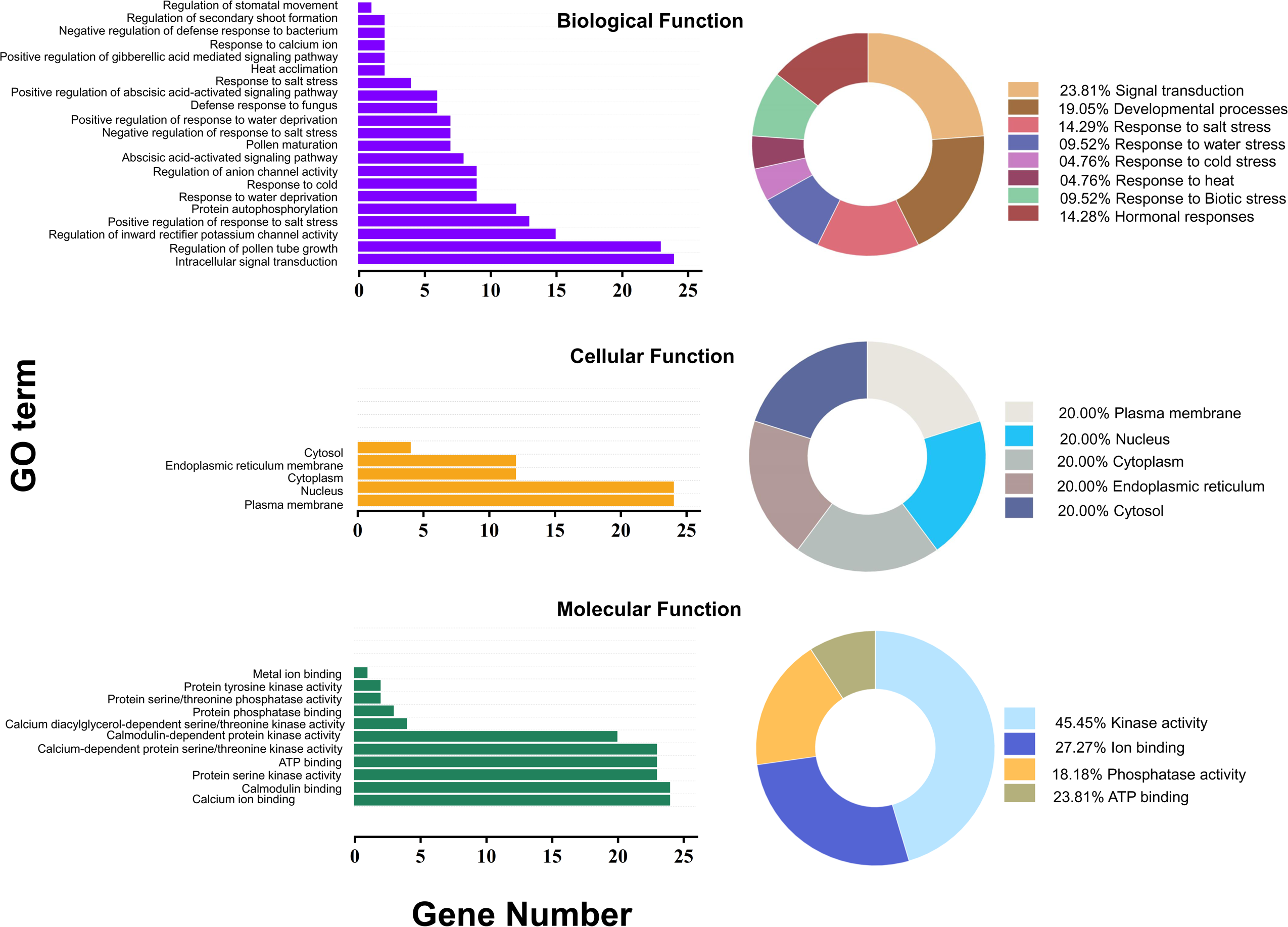
Gene Ontology analysis (GO) annotation was employed to characterize the functional categories of AaCDPK genes. A bar graph presents a summary of gene allocation to various biological processes and responses. Pie charts provide further summaries of the GO annotation results

### 3.7. Protein-protein interaction of AaCDPKs

To investigate the potential regulatory network of AaCDPKs, we constructed an interaction PPI network based on *Arabidopsis* orthologous proteins (**Figure 8**). From the resultant supplementary, we observed that AaCDPKs and other proteins of different pathways interact.

**Fig. 8:**
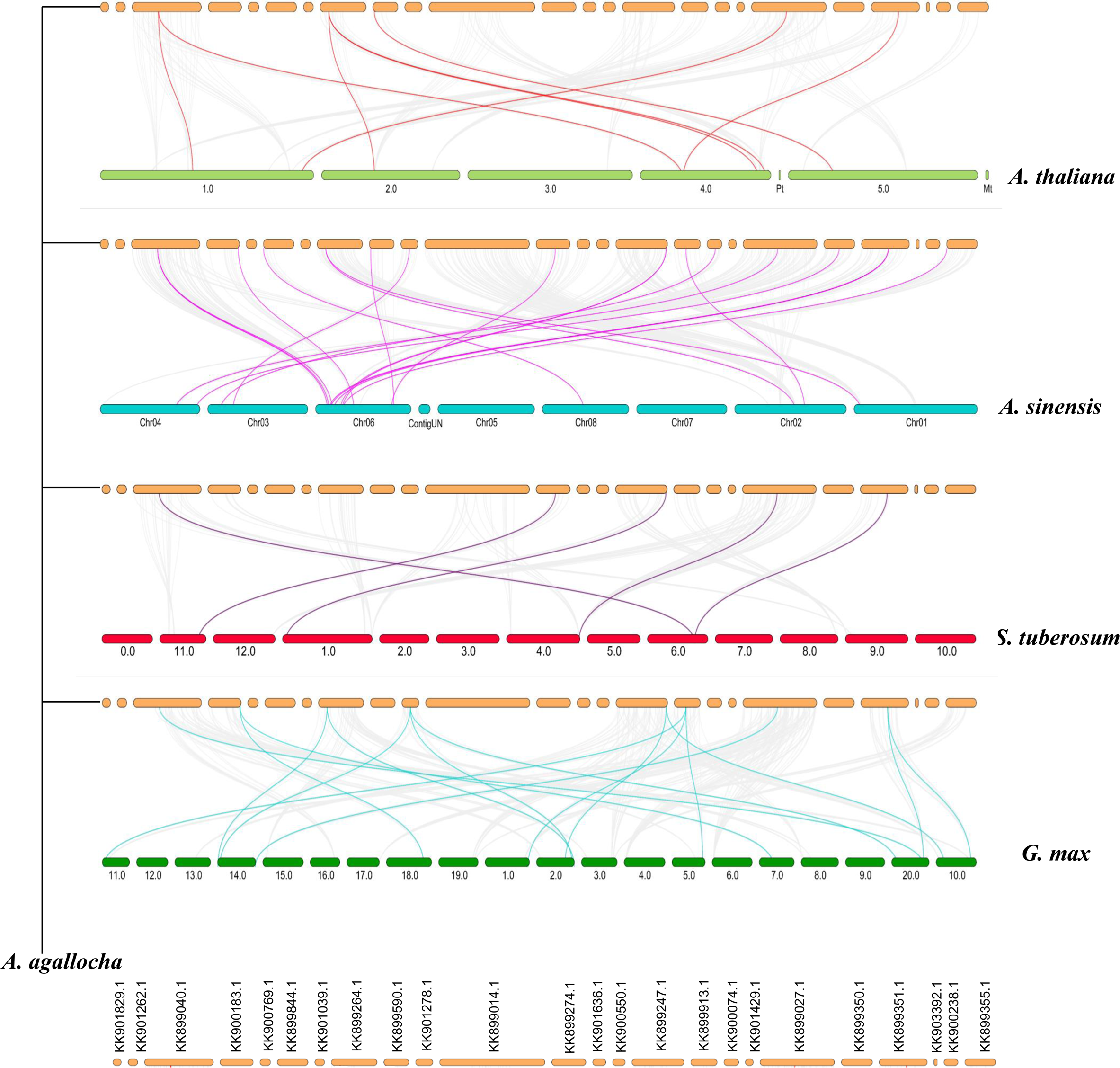
Synteny analysis (A) An overview of the evolutionary relationships of AaCDPK in the genome of A. agallocha. (B). An evolutionary relationship of AaCDPKs was conducted by comparing them to CDPKs of in A. thaliana, G. max and, S. tuberosum. Synteny analysis was employed to identify conserved gene blocks across these angiosperms. Grey lines in the figure represent collinear blocks, while colored lines indicate syntenic CDPK gene pairs.

CDPKs were found to be involved in signalling pathways related to plant growth and development and abiotic stress responses. This analysis showed that AaCPK2 is associated with several respiratory oxidase homolog proteins (RBOH), which promote ROS production. As reported in our previous research, the AaRboh proteins act as signaling molecules for agarwood formation (Begum et al., 2024). It can be stated that AaCDPK10 and AaCDPK11.2 might be involved with drought resistance because of their interaction with protein DEHYDRATION-INDUCED 19 homolog 2 (DI9). Interactions between AaCDPK3, AaCDPK6.2, AaCDPK8, and AaCDPK21 with SLAH3 suggest that these putative AaCDPKs may be involved as negative regulators of guard cell potassium channels. While the aforementioned putative AaCDPKs (apart from AaCDPK3) may be essential for stomatal closure due to their interaction with SLAC1. The interaction between AaCDPK3, AaCDPK6.2, and AaCDPK2 and GORK implies that they may be involved in stomata movement in response to variations in plasma membrane voltage. The PPI investigation suggests that AaCDPKs will likely play essential roles in plant growth, development, and stress responses, including stomatal movement, ion transport, and hormone signalling pathways through ROS production.

### 3.8. Synteny and duplication analysis

Analysis of AaCDPK gene duplication modes, performed using DupGen Finder, identified two specific duplication mechanisms: dispersed duplication and whole-genome duplication. Whole-genome duplication (WGD) resulted in the forming of the *AaCDPK6.1*-*AaCDPK6.2* gene pair, indicative of a duplication event within group 1. Conversely, dispersed duplication events led to the generation of three gene pairs: AaCDPK17.1-AaCDPk17.2, *AaCDPK17.1*-*AaCDPK3*, and *AaCDPK24.1*-*AaCDPK13.1*, representing duplication events within groups 2 and 3. Analysis of Ka/Ks ratios revealed that four duplicated gene pairs of *A*. *agallocha* exhibit values less than 1, indicating they are subject to purifying selection. Divergence times for these duplicate members ranged from 17.97 to 199.27 million years ago (MYA). (Supplementary Table 7).

To understand the evolutionary history of *AaCDPK* genes, we analyzed their syntenic relationships of the genome of *A*. *agallocha* with other plant species genomes, including *A. thaliana*, *A*. *sinensis*, *G*. *max*, *S*. *tuberosum*, and *V*. *vinifera*. We detected eight homologous gene pairs between *A*. *agallocha* and *A*. *thaliana*. The orthologous genes pairs between these two genome are as follows: *AaCDPK11.2- AtCDPK11*, *AaCDPK11.2*- *AtCDPK4*, *AaCDPK6.1*- *AtCDPK6*, *AaCDPK6.1*- AtCDPK26, *AaCDPK6.1*- *AtCDPK5, AaCDPK17.1*- *AtCDPK34*, *AaCDPK29*- *AtCDPK29* and *AaCDPK11.1*- *AtCDPK4*.Comparing the genomes of *A. sinensis*, *G*. *max*, *S*. *tuberosum*, and *V*. *vinifera* to that of *A*. *agallocha*, we found twenty, sixteen, five, and ten pairs of homologous CDPKs genes, respectively (**Figure 9, Supplementary Table S8**).

**Fig. 9:**
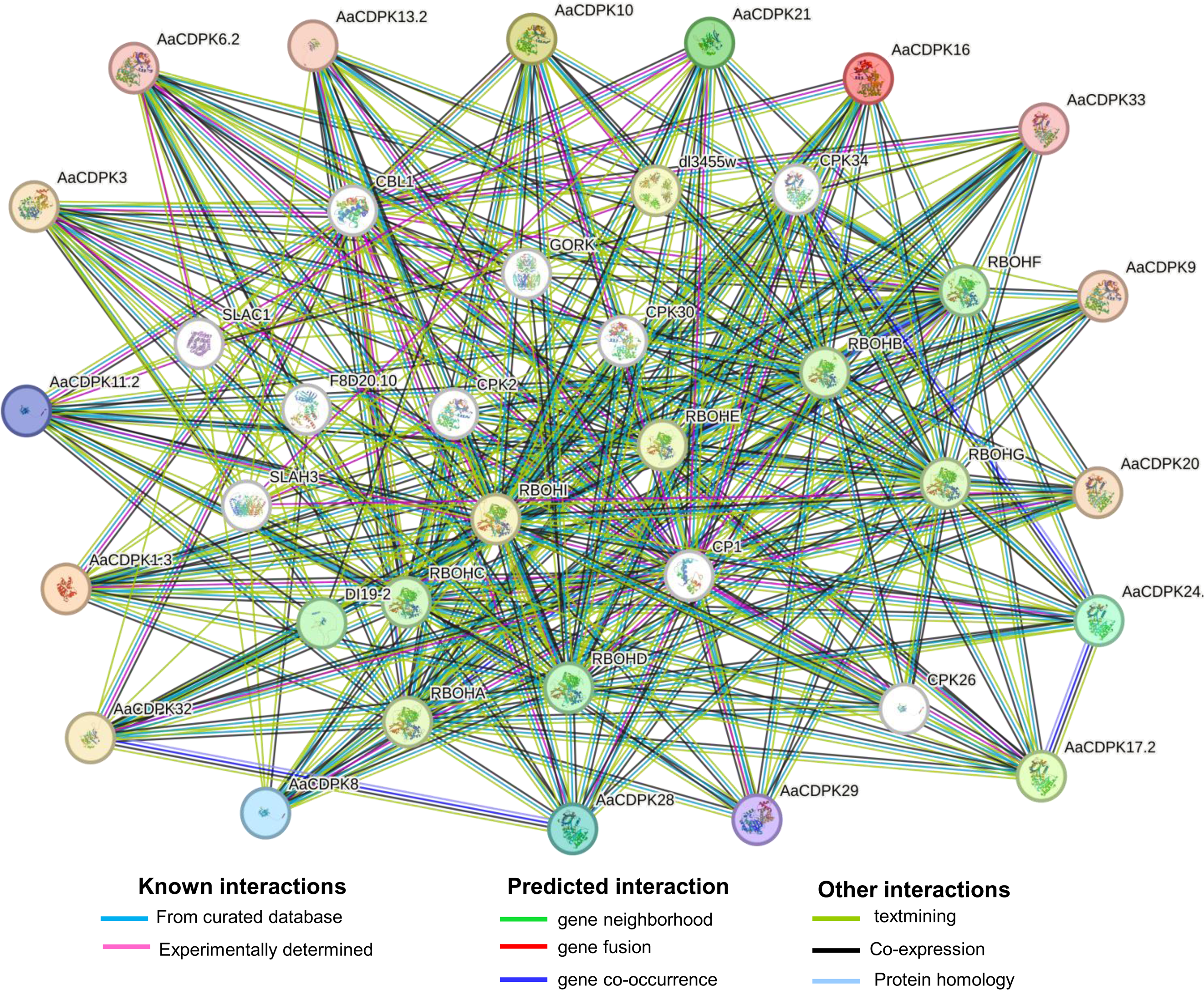
A. agallocha’s AaCDPK protein interaction network is based on orthologues from Arabidopsis. A network model of proteins shows the possible AaCDPK and their functional partners in each enriched route. The lines of different colors show the kinds of interactions that may occur between the putative AaCDPK and their functional partners.

### 3.9. Protein modeling, model quality assessment, and validation

Homology modeling was performed for all AaCDPK isoforms to investigate their tertiary structures. The tertiary structures of the AaCDPK proteins are primarily composed of three secondary structural elements: α-helices, β-strands, and coils. (**Figure 10**). Homology modeling revealed that the 3D structures of AaCDPK proteins primarily consist of α-helices, with coils making up a much smaller proportion. The high proportion of α-helices as secondary structures indicates the stability of the proteins. In all AaCDPK proteins, structural folds composed of green and purple α-helices and β-strands represent the catalytic kinase domain. The other important motif, EF-hand, comprises two secondary structures: α-helices and coils. The template ID, MolProbity Score, Clash Score, and Ramachandran favored percentage of the homology-modeled structures of putative AaCDPKs are listed in Supplementary Table S8. The MolProbity score and clash score of the modeled proteins ranged from 1.31 to 1.98 and 1.31 to 5.46, respectively, indicating good quality of models. Additionally, all protein model structures exhibited a Ramachandran favored percentage above 90.28%. The quality factor of the models, as determined by ERRAT, varied between 92.04 and 96.92, suggesting the acceptable quality of the built models (**Supplementary Table 9**).

**Fig. 10:**
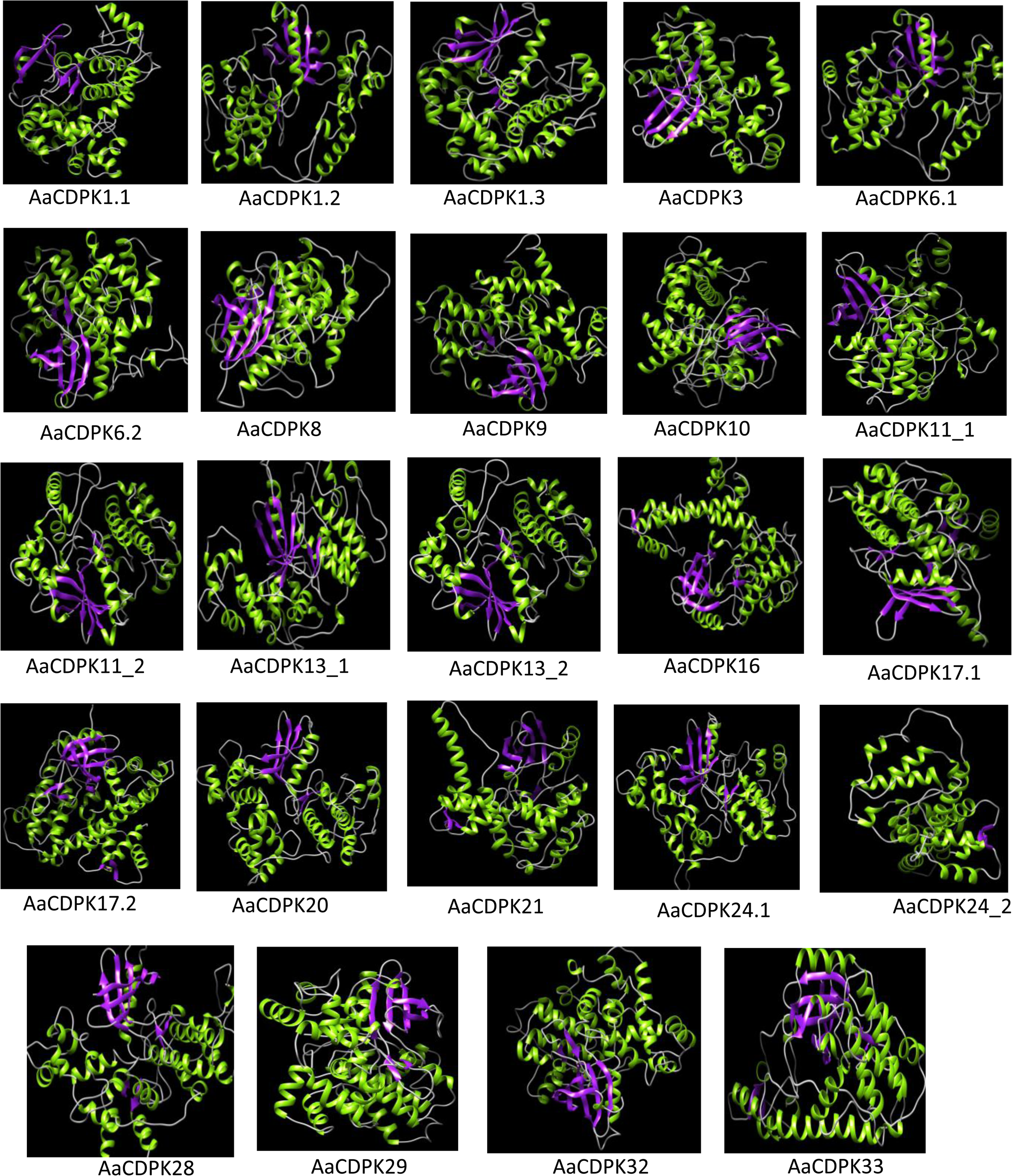
3D protein structure prediction for 24 AaCDPK proteins was performed using UCSF CHIMERA 1.10. Different colors were used to visualize the predicted protein structures, highlighting potential strand, helical, and coil regions.

### 3.10. Expression pattern of candidate AaCDPKs under the treatments of MeJA, H_2_O_2,_ and CaCl_2_

Methyl jasmonate (MeJA), hydrogen peroxide (H_2_O_2_), and calcium chloride (CaCl_2_) are well-established as pivotal signaling molecules in the biosynthetic pathway of agarwood deposition (Xu et al., 2016; Lv et al., 2019). Furthermore, exogenous CaCl_2_ has been shown to induce CDPK gene expression (Liu et al., 2006). Based on these observations, quantitative real-time PCR (qRT-PCR) analysis was conducted to examine the expression patterns of AaCDPK genes in calli in response to separate treatments with MeJA, H_2_O_2_, and CaCl_2_.Ten AaCDPK genes (*AaCDPK* 1.1/6.2/10/11.1/11.2/13.2/17.2/20/21 and 32) were selected for expression analysis from a set of putative AaCDPK genes. These genes were chosen based on their distinctive RNA-seq expression profiles in agarwood tissue and their promoter characteristics (**Figures 5 and 6).** All treatments resulted in a significant upregulation of six AaCDPK genes, while the expression of four genes remained relatively unchanged (**Figure 11**). Notably, *AaCDPK32*, *AaCDPK10*, and *AaCDPK17.2* exhibited the most differential upregulation. MeJA treatment significantly upregulated the expression of AaCDPK genes in calli. Specifically, *AaCDPK10* expression peaked at 6 hours with a 14.84-fold increase, while *AaCDPK32*, *AaCDPK1.1*, *AaCDPK21*, *AaCDPK17.2*, and *AaCDPK20* peaked at 6 hours with 12.32-fold, 7.03-fold, 4.81-fold, 2.499 fold and 2.90-fold increases, respectively. Subsequently, the expression of these genes declined from 6 to 48 hours (except for AaCDPK20 and AaCDPK21). Remarkably, AaCDPK20 and AaCDPK21 exhibited a similar fold-change in expression from 6 to 8 hours. Notably, no significant changes in the expression levels of AaCDPK6.2, AaCDPK11.1, AaCDPK11.2, and AaCDPK13.2 were observed in any of the three treatments compared to the control calli across all time points.

**Fig. 11:**
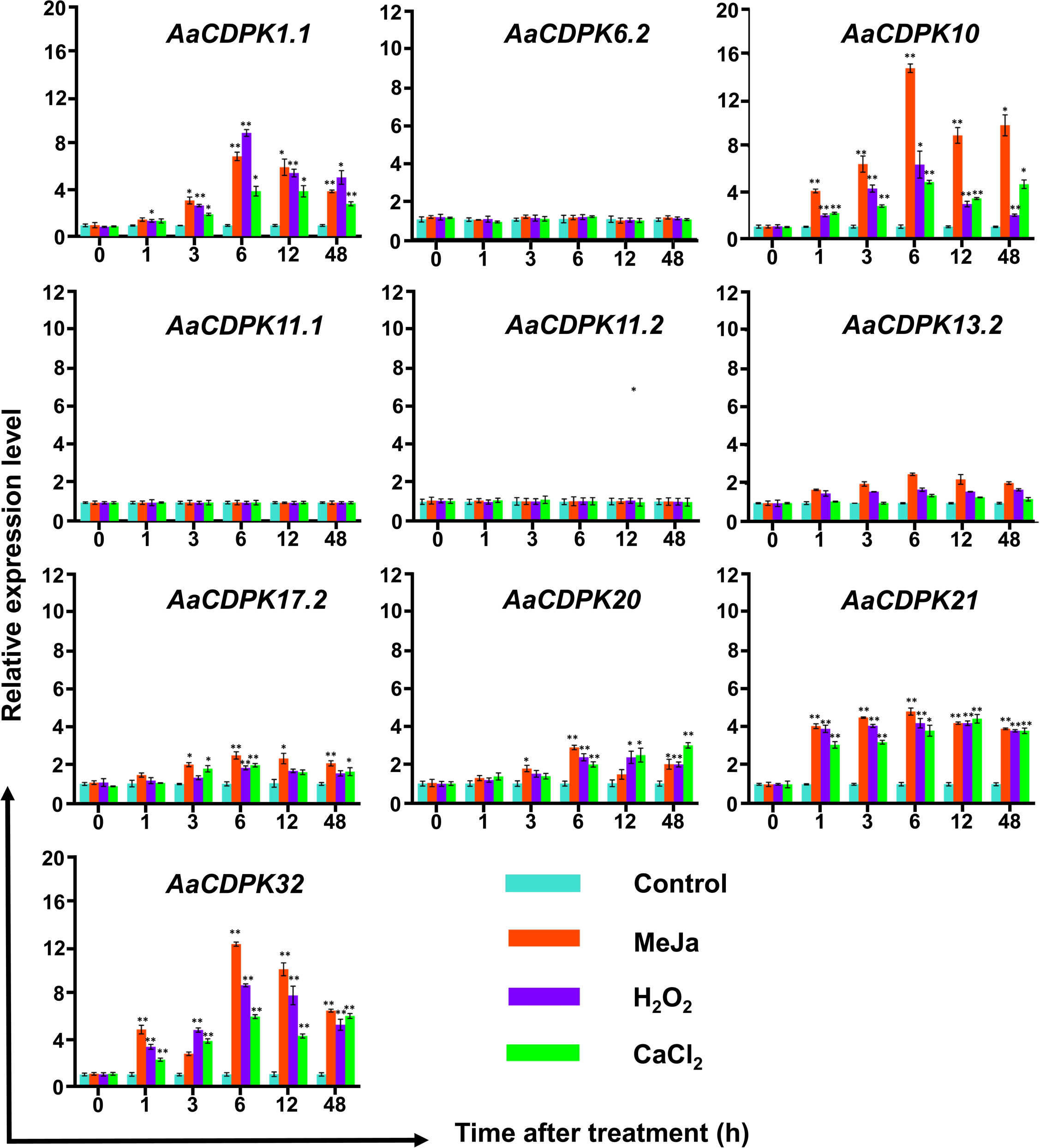
Relative expression levels of AaCDPK genes in treated calli. relative transcripts abundance of 10 selected AaCDPK were measured in calli tissue transferred to MS media with Meja, H2O2 and CaCl_2_ respectively and calli without any treatment considered as control condition and samples harvested at 0,1,2,6,12,24 and 48 h. Transcript abundances were measured using A. agallocha GAPDH as internal control. Asterisks (*) denotes a significant difference compared with healthy samples at *P< 0.05 or **P< 0.01 (Student’s t-test). Data represent means ± SE off three independent experiments).

## Discussion

The role of the CDPK gene family during complex defense mechanisms of A. agallocha has been unraveled. A comprehensive analysis of the CDPK multi-gene family was carried out, and 24 *AaCDPK* genes were identified in the genome of *A*. *agallocha*.

### 4.1 Characteristics of the *CDPK* gene family in *A*. *agallocha*

The biochemical nature of the identified AaCDPK proteins is primarily acidic, with isoelectric points (pI) ranging from 5 to 7. However, putative AaCDPK18 and AaCDPK28 from group IV and AaCDPK10 from Group III had pI values above 9 and 7, respectively. The results obtained in this study align with the previously reported work (Nuruzzaman et al., 2010; Hamel et al., 2014; Cai et al., 2015; Zhang et al., 2017; Zhang et al., 2020). Plant CDPKs are acidic due to serine/threonine residues in the conserved protein kinase domain and acidic residues in the activation loop. These acidic residues are proposed to prevent premature phosphorylation, maintaining the kinase’s inactive state until activation of CDPK proteins (Reinhardt and Leonard., 2023). The domain confirmation search for the identified AaCDPK proteins is characterized by the presence of conserved protein kinase (PF00069) and EF-hand (PF13499) domains (Asano et al., 2005; Kong et al., 2013; Mu et al., 2024). The kinase domain phosphorylates specific amino acids in target proteins, altering their conformation and function (Hank et al., 1988), and EF-hand domains endow the protein with the ability to perceive Ca^2+^ signals into diverse biochemical responses (Dekomah et al., 2022). The AaCDPK proteins identified in this study have two to four EF-hand domains. Motif analysis has shown that all the putative AaCDPKs’ protein kinase domains possess ATP binding sites (GO:0005524) (**Figure 4 and Supplementary Table 4**), harnessing ATP to phosphorylate target proteins. The phosphorylation activity of AaCDPK proteins suggests their role in regulating signal transduction pathways in *A*. *agallocha* (Wen et al., 2020; Li et al., 2024).

Bioinformatic analysis revealed that nine of the 24 AaCDPKs possess an N-myristoylation motif, characterized by a crucial glycine residue at the N-terminus. Myristoylation was thought to be essential for protein-protein and protein-lipid interactions, facilitating membrane localization and involvement in diverse signal transduction cascades (Johnson et al., 1994; Farazi et al., 2001). CDPKs, crucial Ca^2+^ sensors, often possess N-myristoylation motifs that target CDPK proteins to the plasma membrane, suggesting a role for myristoylation in membrane binding of protein and Ca^2+^ signal transduction (Lu, S. X., & Hrabak, E. M., 2013, Mehlmer et al., 2010). All the AaCDPKs identified in this study have palmitoylation sites characterized by Cys residue at positions 3, 4, or 5, indicating the possibility of further membrane anchoring through palmitoylation. . Previous research has demonstrated that N-myristoylation and palmitoylation sites are key in determining CDPK subcellular localization (Zhao et al., 2021a). Furthermore, predicted subcellular localization suggests that AaCDPK proteins are predominantly found in cytoplasm, chloroplasts, and mitochondria.

### 4.2. Evolutionary relationships and collinearity analysis of AaCDPKs

Based on a phylogenetic analysis of the AaCDPK and AtCDPK, the proteins are clustered into four distinct groups. This classification is congruent with previous phylogenetic studies in cucumber (Xu et al., 2015), grape (Zhang et al., 2015); pineapple (Zhang et al., 2020); sea- island cotton (Shi., G.& Zhu., X., 2022); and Populus (Ma et al., 2024). This classification suggests conserved evolutionary patterns within the *CDPK* gene family across diverse plant species. The conserved gene structure and motif distribution among members of the same phylogenetic Group of AaCDPK proteins (**Figure 2A, B**) is consistent with the CDPK structural features in *Capsicum annuum* (Cai et al., 2015), *Hordeum vulgare* L (Yang et al., 2017), and *Avena sativa* L (Li et al., 2024). This further supports their close evolutionary relationships and phylogenetic classification. D distinct gene structure and motif composition variations were observed among the four phylogenetic groups. These differences suggest potential functional divergence among the groups.

The *CDPK* gene family has significantly expanded over evolutionary time, increasing from approximately four genes in the ancestral land plant to around 30-40 genes in angiosperms via different duplication mechanisms. (Valmonte et al., 2014, Gardette et al., 2014, Dokemah et al., 2022). One pair of the four duplicated *AaCDPK* genes (*AaCDPK6.1*/*AaCDPK6.2*) was duplicated through WGD and the remaining three pairs (*AaCDPK17.1*/*AaCDPK17.2*, *AaCDPK17.1*/*AaCDPK3*, and *AaCDPK24.1*/*AaCDPK13.1*) duplicated through dispersed gene duplication mechanism. Among different duplication mechanisms, WGD and dispersed. The frequent WGD of CDPK genes, encoding essential calcium sensors in plant stress response pathways, confers a significant evolutionary advantage. This duplication facilitates functional diversification, enabling adaptation to a broad spectrum of environmental stresses by evolving multiple gene copies with potentially specialized functionalities (Mu et al., 2024). The Ka/Ks values of duplicated gene pairs were less than 1, indicating they undergo purifying selection (Zou et.al., 2024). This suggests that natural selection maintains the amino acid sequence, which is crucial for the protein’s functionality. The purifying selection, which is sometimes referred to as negative selection, works to get rid of harmful mutations that could impair the function of proteins (Hartl and Clark, 2007)

Identifying genes with shared ancestry (orthologs) across species is vital for functional genomics, as it helps to gain a more comprehensive understanding of the evolutionary history of genes and plants (Ming et al. 2013, Yan et al., 2024). Synteny analysis assessed the phylogenetic relationships between *AaCDPK* genes and their homologs genes in four plant species: *A*. *thaliana*, *A*. *sinensis*, *S*. *tuberosum*, and *G*. *max*. The analysis identified the most orthologous genes between *A*. *agallocha* and *A*. *sinensis*. This orthology supports their close evolutionary affinity and belonging to the same genus concerning their gene composition. Based on orthologous gene counts, the next most closely related plant species were *G*. *max*, *A*. *thaliana*, and *S*. *tuberosum*. These orthologous gene pairs may have originated from a last common ancestor (Gabaldon and Koonin 2013). Therefore, observing more orthologous gene pairs in two plant species allows us to infer a closer evolutionary relationship. Orthologous gene pairs are generally assumed to retain equivalent functions in different species (Steenwyk et al., 2022).

### 4.3. AaCDPKs are key players in diverse biological processes, especially stress responses

The different types of CRES present in the promoter can provide insights into the genes’ potential functions and regulatory mechanisms (Li et al., 2021). The presence of LTR, WUNmotif, ARE, WRE3, TC-rich repeats, etc., in the promoter regions of the putative *AaCDPK*s suggests that they play a pivotal role when *Aquilaria* plants encounter various biotic and abiotic stresses. The functional analysis of the AaCDPKs also supports the role of AaCDPK during stresses. Stress-responsive CRES was also reported in the promoters of *CDPK* genes of numerous plant species, including tomato (Li et al. 2022; potato (Dekomah 2022), and patchouli (Liu et al. 2023). Furthermore, the promoters of nine putative *CDPK*s harbor CRES elements, which are known to be involved in plant developmental processes. Prior research has demonstrated that the phytohormone methyl jasmonate plays a significant role in triggering the production of sesquiterpenes in *Aquilaria* species as a defense mechanism against diverse stressors (Xu et al., 2013; Xu et al., 2016; Lv et al., 2019, Naziz et al. 2019, Faizal et al. 2021). The TGACG-motif and CGTCA-motif were identified in the putative AaCDPK gene promoters, which are the two MeJA-responsive cis-regulatory elements (Nejad et al., 2012; Liu et al., 2022). Interestingly, the promoter regions of all the differentially expressed members (*AaCDPK10*/*32*/*1*/*21*/*20*) in RNA-seq data (except *AaCDPK10*) also possess the above-mentioned MeJA responsive Cis-element. The promoter region of *AaCDPK10*, which did not have either of these motifs, contained Cis-element for ethylene and salicylic acid phytohormone. It is a fact that three phytohormones, i.e., MeJA, Ethylene, and Salicylic acid, are responsible for sesquiterpenes accumulation for agarwood formation (Liu et al., 2021). Combining analysis of result analysis of CRES and in-silico expression, it can be inferred that the differentially expressed upregulated putative AaCDPKs may involved with agarwood formation, acting as signalling molecules for sesquiterpene biosynthesis.

*CDPK* genes exhibit rapid and transient differential expression under stress conditions (Boudsocq and Sheen, 2012; Dekomah et al., 2022; Lv et al., 2024). The six *AaCDPK*s (AaCDPK*10*/*32*/*1*/*17.2*/*21*/*20*) were upregulated in calli tissue treated with MeJA, H_2_O_2_, and CaCl_2_, respectively. Previously, differential expression of five *CDPK* genes (*CDPK1*, *CDPK*2, *CDPK3*, *CDPK5*, and *CDPK6*) had been recorded in MeJA-treated calli tissue of *A. sinensis* (Xu et al., 2013). These five *A*. *sinensis* genes show homology with the *AaCDPK* genes studied here as follows: *CDPK1* = *AaCDPK1.1*, *CDPK2* = *AaCDPK32*, *CDPK3* = *AaCDPK11.2*, *CDPK5* = *AaCDPK6.2*, and *CDPK6* = *AaCDPK10*. Three *AaCDPK* genes (*AaCDPK10*/*32*/*1.1*) exhibited similar differential expression patterns in *A*. *agallocha* and *A*. *sinensis*. Moreover, three putative AaCDPKs (*AaCDPK17.2*, *AaCDPK20*, and *CDPK21*) showed differential expression patterns in this study. However, *AaCDPK6.2* and *AaCDPK11.2*, which were differentially expressed in *A*. *sinensis*, showed no expression in *A*. *agallocha*. It can be speculated that upregulated putative *AaCDPK*s, along with their substrate AaRboh proteins, act as signaling molecules to initiate JA, SA, and ET phytohormones, which potentially contribute to agarwood sesquiterpene formation **(Figure 12**) ((Xu et al., 2016; Zhang et al.; 2021; Ma et al., 2023). The role of CDPK proteins during defence mechanisms has been reported in many plant species. Arabidopsis CPKs are essential for plant immunity. AtCPK1 activates NADPH oxidase (Rboh), triggering oxidative burst (Dubiella et al. 2013) and phosphorylates Phenylalanine Ammonia-lyase, inducing salicylic acid accumulation (Coca and Segndo, 2010). NLR-dependent pathogen resistance relies on the phosphorylation of specific WRKY transcription factors (*WRKY8*, *28*, and *48*) by *AtCPK4*, *5*, *6*, and *11*(Shu et al., 2020). Like Arabidopsis, silencing *CDPK2* genes in tobacco resulted in a weakened and delayed hypersensitive response to the fungal Avr9 elicitor (Boudosocq and Sheen, 2012). The induction of *OsCPK10* expression upon treatment with a *Magnaporthe grisea* elicitor highlights the involvement of *CPK*s in rice defense against fungal pathogens (Fu et al., 2013). The activation of *StCDPK7* contributes to various plant defense responses against fungal elicitors (Romeis et al., 2001; Coca and San Segundo, 2010). The *Pseudomonas syringae* pv. tomato DC3000 infection upregulates most tomato *CDPK*s and Phytohormones (ET, JA, SA) involved in biotic stress-induced BrrCDPKs and BrrRbohD1/D2 expression in *Brassica rapa* var. rapa, supporting a role for *CDPK*s in plant defense. (Dekomah et al., 2022). The observed differential expression patterns of *CDPK* genes in *A*. *agallocha,* along with the study PPI, promoter analysis, gene ontology study, and in-silico study, strongly suggest their involvement in defense responses, consistent with well-established roles for *CDPK*s in plant immunity across plant various species.

**Fig. 12:**
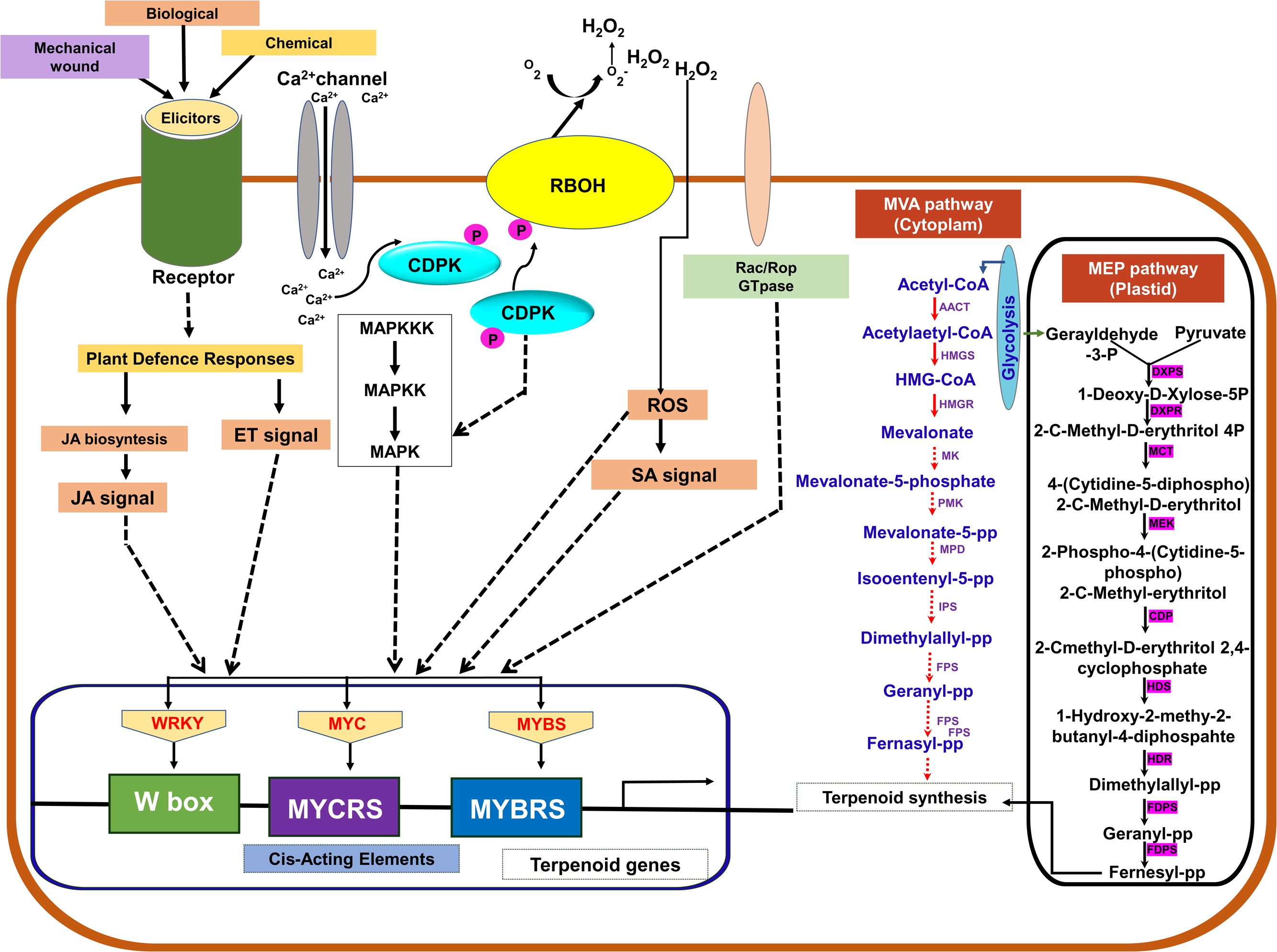
Signal transduction mechanism of sesquiterpene biosynthesis pathway of agarwood resin deposition.

## Conclusion

This study comprehensively analyzed *A*. *agallocha CDPK* genes, encompassing phylogenetic relationships, conserved motifs, gene structures, promoter analysis, in silico expression, gene ontology, PPI, secondary and tertiary structure, and expression profiles. A total of 24 *AaCDPK* genes were identified and classified into four distinct groups. Motif and gene structure analyses revealed conserved patterns within each Group. PPI, Promoter analysis, including in silico predictions and qRT-PCR experiments, strongly suggests the involvement of AaCDPK protein as signaling molecules in stress-related sesquiterpene synthesis pathways. Furthermore, PPI results demonstrate interactions between AaCDPK and Rboh proteins, indicating the interplay of two secondary messengers: calcium and ROS in *A*. *agallocha* defense. The differential expression levels of six *AaCDPK*s observed under different stress treatments suggest that these genes play crucial roles in the signal transduction mechanism of the sesquiterpene biosynthesis pathway in *A*. *agallocha* and contribute to diverse biological functions to provide self-defense against the stresses. Thus, it can be inferred that this gene family likely plays a vital role in transducing stress signals associated with agarwood formation in agarwood-producing species.

## Data availability

The data supporting the finding of this study is provided in the manuscript and its supplementary material.

## Ethics approval

Not applicable

## Acknowledgement

The authors are indebted to Gauhati University for providing the technical facility. The authors also acknowledge Department of Biotechnology (DBT), Government of India for providing the financial aid.

## Abbreviations

AaCDPK: Aquilaria agallocha calcium dependent protein kinase
ROS: Reactive Oxygen Species
HMM: Hidden Markov Model
CDS: Coding Sequences
GO: Gene ontology
KEGG: Kyoto Encyclopedia of Genes and Genomes database

## Supplementary files

**Supplementary figure 1:** Flowchart of methodology

**Supplementary Table 1:** Details of primer used in this study for qRT-PCR analysis

**Supplementary Table 2:** Details of A. agallocha *AaCDPK*s

**Supplementary Table 3**: Annotation of ten conserved motif of AaCDPKs

**Supplementary Table 4:** Details CRES elements of AaCDPK

**Supplementary Table 5**: Log fold value of putative CDPKs of A. agallocha

**Supplementary Table 6:** Duplication genes pair and divergent time

**Supplimentary Table 7:** Orthologous gene pairs

**Supplementary tabel 8**: Quality asssement details of protein structure build with homology modeling

