## SupplementaryImage 1 for "Genome-wide identification and expression analysis of CDPK proteins in agarwood-producing Aquilaria agallocha trees"

### Slide 1
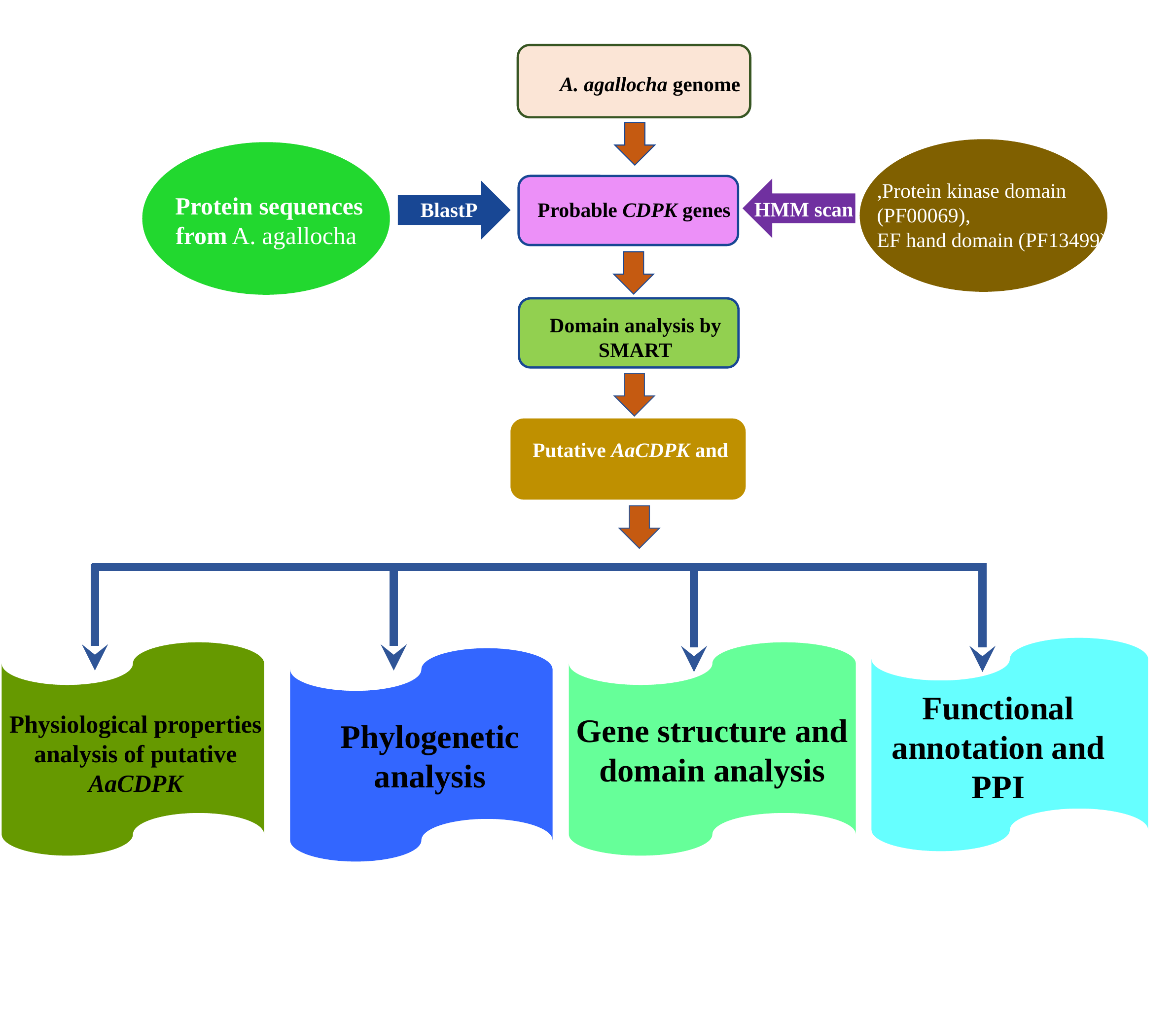

Protein sequences from A. agallocha
HMM scan
BlastP
Probable CDPK genes
Domain analysis by SMART
Putative AaCDPK and
Gene structure and domain analysis
Functional annotation and PPI
Physiological properties analysis of putative AaCDPK
A. agallocha genome
,Protein kinase domain
(PF00069),
EF hand domain (PF13499)
Phylogenetic analysis
